# *Nanoscape*, a data-driven 3D real-time interactive virtual cell environment

**DOI:** 10.1101/2020.10.15.340778

**Authors:** Shereen R. Kadir, Andrew Lilja, Nick Gunn, Campbell Strong, Rowan T. Hughes, Benjamin J. Bailey, James Rae, Robert G. Parton, John McGhee

## Abstract

Knowledge of cellular and structural biology has reached unprecedented levels of detail. In conjunction with advances in 3D computer visualisation techniques this has allowed exploration of cellular ultrastructure and environments by a virtual user. The extraction and integration of relevant scientific information, along with consideration of the best representation of data, is often a bottleneck in the visualisation process for many 3D biomedical artists. Here we introduce ‘Nanoscape’, a collaborative project between 3D computer artists, computer graphics developers, and cell biologists that enables a user to navigate a cell in a complex 3D computer visualised environment. We combine actual data from various scientific disciplines (including structural biology, cell biology and multiple microscopic techniques) and apply artistic expression and design aesthetics to create a unique new experience where a real-time virtual explorer can traverse a cell surface, observe and interact with a more scientifically accurate cell surface environment.

## Introduction

Since cells were first observed under the microscope in the 17^th^ Century, advances in experimental science and technology have greatly improved our understanding of cell biology (1). Modern imaging techniques have played a central role in unravelling the intricacies of the cellular landscape at the molecular level. Atomic- or near-atomic-resolution views of individual proteins, macromolecular assemblies and cellular architecture can be achieved though methods such as X-ray crystallography, NMR spectroscopy, and cryo-electron microscopy. Live cell imaging enables scientists to observe cellular structures, processes, and behaviour in real-time, and in recent decades significant enhancements in fluorescent tags and biosensors have shed considerable light on protein kinetics, interactions, and diffusion (1, 2).

3D visualisation of scientific concepts and experimental data is becoming an increasingly popular communication tool that can provide insight where traditional 2D graphical illustration or descriptive text cannot (3). The educational benefits are numerous as they can clarify complex or abstract concepts (4–6), and exploratory visualisations may help test hypotheses and generate new ideas (7).

Since no single experimental modality is sufficient to elucidate the structure and dynamics of macromolecular assemblies and cellular processes, integrative modeling of data from multiple complementary experimental techniques and scientific disciplines such as molecular biology, biochemistry, biophysics, and computational biology is crucial (8). Despite a wealth of information in scientific journals and bioinformatics databases that are widely available for researchers, careful extraction and interpretation of relevant data, along with consideration of its representation is a time-consuming hurdle for most 3D biomedical visualisers. The nature of multifaceted scientific resources makes it hard for non-scientists to identify these data easily, and there is an urgent need to make platforms more accessible to the biomedical visualisation community (9).

Another long-standing issue in the biomedical visualisation field is the lack of established systems of citation once a molecular animation has been created, and often viewers are left to assume the creators have researched the topic thoroughly. More transparent indications of whether the visualisations have been informed by empirical data along with to links those original sources; highlighting exploratory hypotheses and where speculation or artistic license has been used will give credence (10, 11).

The 3D computer animation and modelling software such as Autodesk Maya (12), SideFX Houdini (13) and Pixologic Zbrush (14) often used by the games and entertainment industry are now widely used by biomedical animators. The true complexity of cellular environments is however often deliberately diminished due to various technical limitations of computer graphics (CG), project time constraints and capabilities of the 3D software operator(s), but also to clarify or emphasise particular features and mechanisms of interest (10). The last couple of decades have seen an increase in visualisation software tools and accurately scaled complex static reconstructions at the molecular level. Notable examples are the HIV-1 virus and *Mycoplasma mycoides* using the packing algorithm CellPACK (15, 16), and a snapshot of a synaptic bouton (17) based on integration of a plethora of imaging techniques, quantitative immunoblotting, and mass spectrometry. Some groups have incorporated simulations of Brownian dynamics or molecular dynamics in 3D atomic resolution models of bacterial cytoplasmic subsections (*Escherichia coli* (18) and *Mycoplasma genitalium* (19, 20)) to examine the effects of protein interactions, stability, and diffusion under crowded cellular environments. Furthermore, mathematical and computational modelling platforms such as V-Cell (21), M-cell (22, 23), and E-cell (24) are designed to be user-friendly for experimental biologists and theoretical biophysicists to run simulations of cell-biological phenomena constrained by a combination of multiple “omics” technologies and imaging data, thereby enabling scientists to analyse 3D representations of their raw data and test hypotheses with varying degrees of molecular and spatial resolution.

Currently many of the existing visualisations and methodologies described above do not fully represent cellular environments due to sheer complexity, issues with gaps in scientific knowledge, computational limitations, and project management concerns including funding and personnel. Our previous work “Journey to the Centre of the Cell (JTCC)”, an immersive Virtual Reality (VR) educational experience of an entire 3D cell based on serial block-face scanning electron microscope (SBEM) imaging data, showed significant improvement in students’ comprehension of cellular structures and processes (25). Although JTCC successfully depicted cellular features from real microscopy data, portrayal of the cell surface environment needed to be oversimplified due to various constraints which included working at higher VR frame rates (90fps +) and a smaller development team. Following on from JTCC, we present “Nanoscape”, a collaborative project between 3D artists, CG developers, and cell biologists to create a first-person interactive real-time open-world experience that enables a user to navigate a breast cancer cell terrain within a tumour microenvironment. This paper shows the integration of data from various scientific disciplines (including structural biology (atomic structures of surface proteins), cell biology and multiple microscopic techniques (such as light microscopy and electron microscopy)) along with artistic approaches taken to more authentically depict molecular crowding, scales, and spatiotemporal interactions in real-time, whilst balancing design aesthetics and the user experience. We present some of the challenges and limitations experienced during the data collection process, conceptualisation of 3D assets and an examination into the practicability of replicating experimental data. We discuss the implications of gaps in scientific knowledge, modification or simplification of data and use of artistic license for visual clarity. It raises important questions about whether molecular visualisations can support outcomes beyond the educational field and help experimentalists to better understand their data.

## Materials and Methods

### Proteins

Protein structures were retrieved from the RSCB protein data bank (PDB) and MoA animations were simulated using the mMaya modelling and rigging kits (Clarafi). Rigged surface or backbone meshes were extracted, animation playblasts recorded in Maya (Autodesk) and composited in After Effects (Adobe CC). See Supplementary Table 1, Supplementary Figure 1 and Supplementary Methods for detail.

For the creation of stylised proteins, backbone meshes extracted from PDB structures were sculpted in Zbrush (Pixologic), polypainted and subsequently texture, normal, and displacement maps were exported. Images were rendered in Maya using Arnold at 1400×1400.

Receptor density simulations on 1um^2^ surface areas (sphere and plane) based on MDA-MB-231 from flow cytometry data (26) were created in Blender 2.78 (Blender Foundation) using the plugin autoPACK (autopack.org) with the spheresBHT packing method. Low poly PDB meshes of CD44, EGFR, EpCAM, Her2, ICAM1 and αVβ3 integrin were created in mMaya.

### Lipids

A cancer lipid bilayer consisting of 400 lipids was simulated using the CHARMM-GUI Membrane Builder based on data from Table 2 Shahane et al. 2019. UCSF Chimera was used to export lipid meshes to Maya.

### Cells, cellular processes and ECM models

Information on dimensions and temporal dynamics were taken from the literature and data from our collaborators (See Supplementary Table 2). All 3D assets were modelled in Zbrush or Maya. Cellular processes were animated in Maya. See Supplementary Methods for further detail.

### Image rendering

Unless stated all images were rendered in Maya using Arnold at 4K (3840×2160).

### Nanoscape open-world compilation

The Unity3D game engine was used to assemble the various components into a coherent representation of a cell surface environment. Static assets were imported into the engine by standard methods. The animated cellular processes and horde of proteins were integrated and simulated in Houdini and output as custom caches that are streamed into the Unity3D engine at runtime.

## Results and Discussion

As part of the pre-production stage, information on the major surface components, processes, and extracellular features commonly found in breast cancer scenarios was collated through a comprehensive review of the literature along with analysis of experimental data obtained from scientific collaborators (Figure 1).

**Figure 1.**
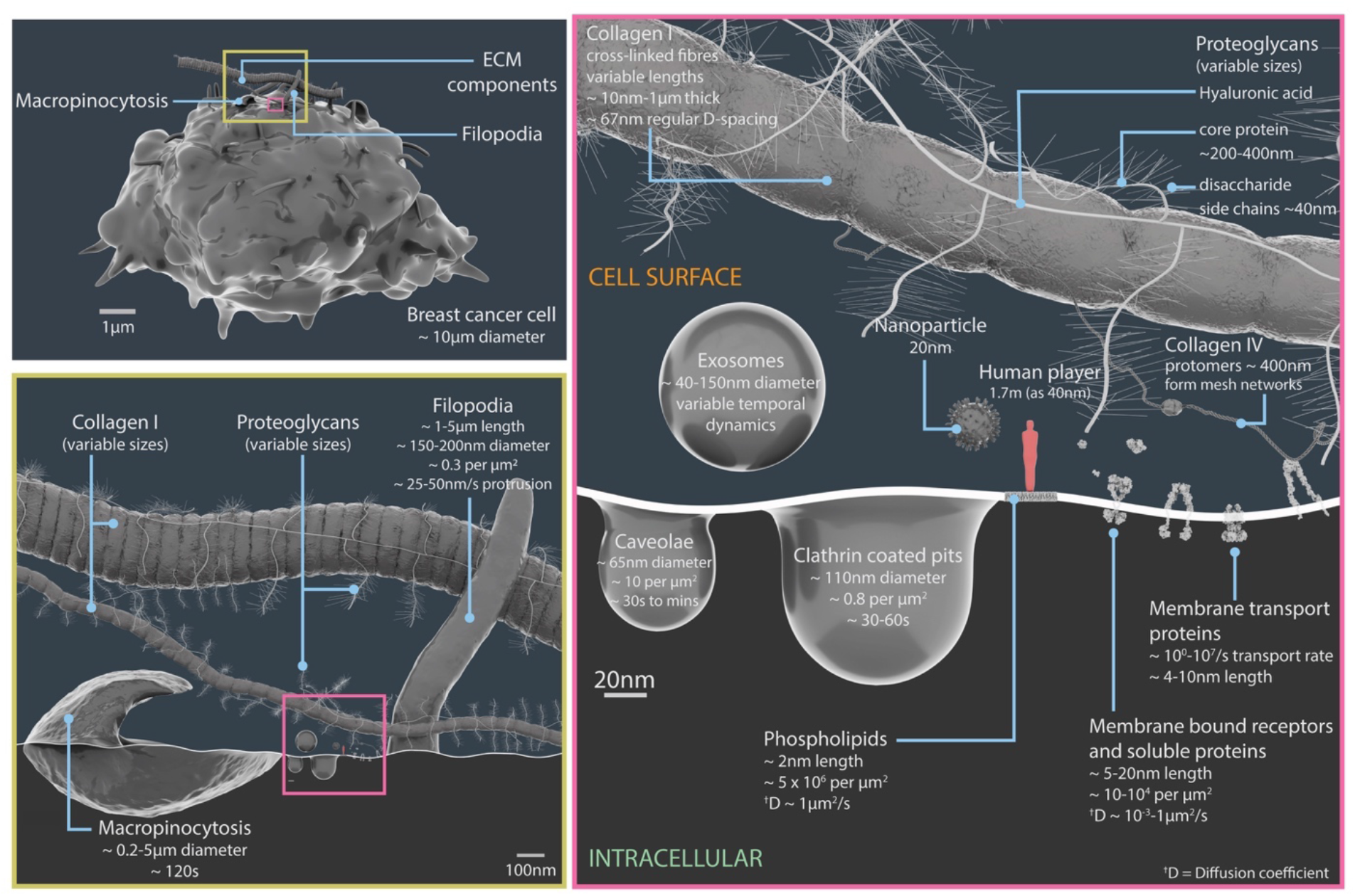
Relative scales and temporal dynamics of cell surface components featured in Nanoscape. A 3D model breast cancer cell with surface structures proportional to a player (light red) standing at an equivalent height of 40nm within the game; yellow insert details ECM components (collagen I and proteoglycans), filopodia and micropinocytosis structures; pink insert details exosomes, caveolae, clathrin coated pits, plasma membrane lipids, surface proteins, ECM components (proteoglycans, collagens I and IV), and a 20nm nanoparticle.

### Cell surface proteins

Cell surface proteins, collectively known as the “surfaceome”, exhibit a wide range of functions, including playing a vital role in communication between the cell and its environment, signal transduction, and transport (27).

The Cell Surface Protein Atlas (CSPA; wlab.ethz.ch/cspa) (28) and the *in silico* human surfaceome (wlab.ethz.ch/surfaceome) (27) which have classified 2,886 entries, were used to first identify different types of surface proteins, for example receptors, soluble, and membrane transport proteins. Subsequently, over 30 prevalent breast cancer-associated surface proteins with structures available from the RSCB protein data bank (PDB) (29) were selected and organised in a cast of characters (Supplementary Table 1 and Figure 2). Where possible, any available information on their motion (molecular dynamics, conformational changes and protein interactions), and population densities were interpreted from various published sources.

**Figure 2.**
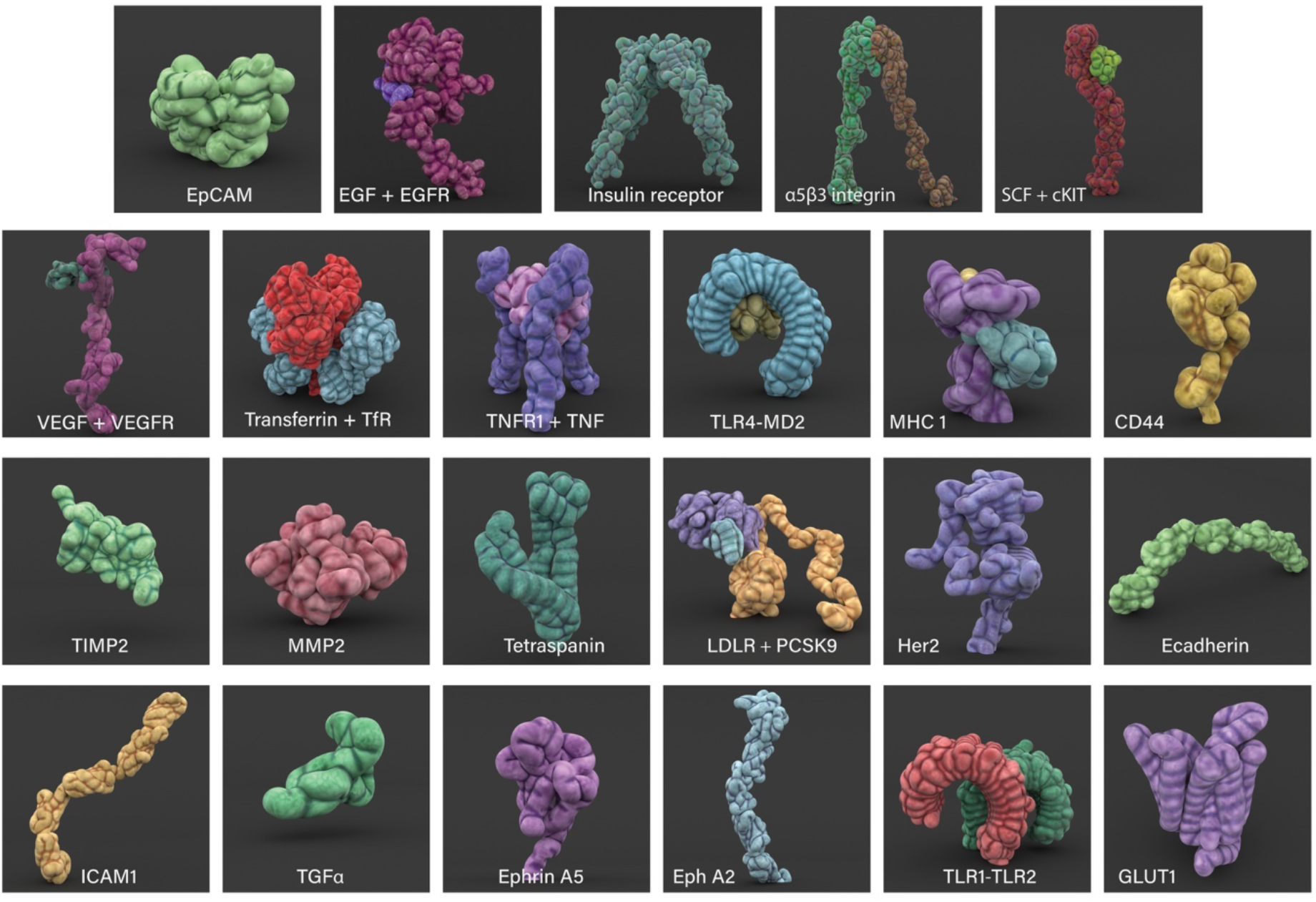
3D modelled cell surface receptors and ligands featured in Nanoscape. Stylised 3D meshes modelled from structures retrieved from the PDB (see Supplementary Table 1 for details). Most proteins are depicted as monomers.

Figure 3A summarises structural and dynamic information gathered about the ErbB family of proteins (which consists of 4 receptor tyrosine kinases: EGFR, Her2, Her3, and Her4) and the associated literature references used to create mechanism of action (MoA) animations for each family member (Supplementary Figure 1A). mMaya modelling and rigging kits (30) which qualitatively replicate molecular dynamics, were used to simulate ligand binding events and transitions between conformational states (see Supplementary Methods). These rudimentary MoA animations provided an interpretation of protein movement based on experimental data available and were subsequently used to inform the artistic design team how best to approach rigging a refined, stylised 3D protein mesh (Figure 3B) using traditional rigging methods.

**Figure 3.**
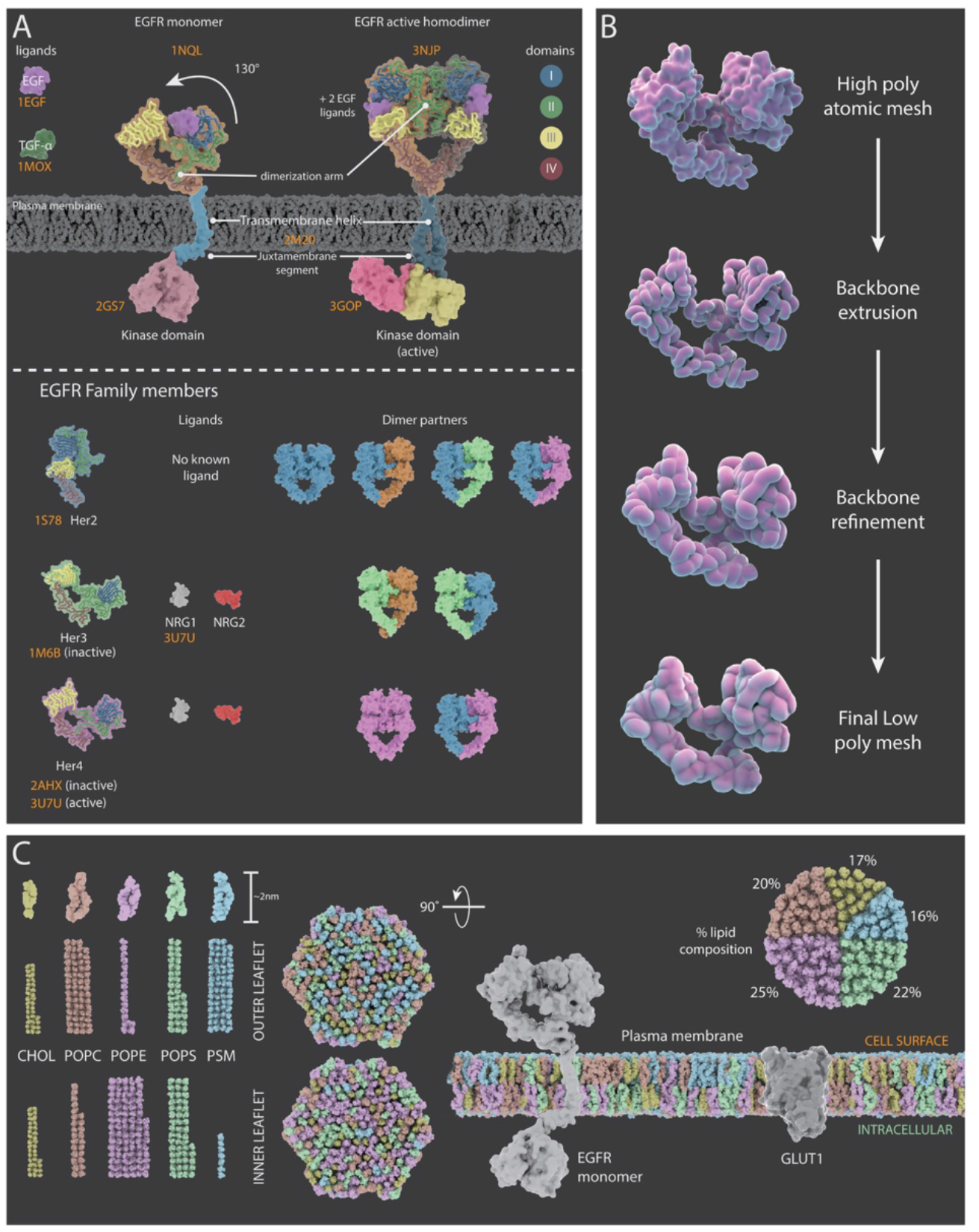
Representations of surface receptors and plasma membrane lipids. (A) Summary of structural and dynamic information collected about the ErbB family of proteins for creation mechanism of action animations. PDB structures meshes (orange text) for ErbB family members (EGFR, Her2, Her3, and Her4) and their ligands. Top: mechanism of action for EFGR, which upon ligand binding the inactive monomer undergoes a conformational change (130° movement), into the active extended conformation and dimerization with another active EGFR protein can occur. Bottom: Her2, Her 3 and Her 4 PDB structures and dimer partner combinations. (B) Creation of stylised protein meshes. Original high poly atomic structures sourced from the PBD with their backbones extruded and refined to produce a low poly stylised mesh. (C) A 3D model composition of a representative cancer lipid bilayer highlighting an asymmetric distribution of 400 lipids (data adapted from (31)). Bilayer components include cholesterol (CHOL), 1-palmitoyl-2-oleoyl-sn-glycero-3-phosphocholine (POPC), 1-palmatoyl-2-oleoyl-sn-glycero-3-phosphoethanolamine (POPE), 1-palmitoyl-2-oleoyl-sn-glycero-3-phospho-L-serine (POPS), and palmitoylsphingomyelin (PSM). Bar charts plot the proportion of each lipid species within the outer and inner leaflets; pie chart represents the percentage lipid compositions within the bilayer. Side view illustration of a model cancer plasma membrane with proteins EGFR and GLUT1.

Conformational flexibility plays a crucial role in enabling protein-ligand interactions, multi-specificity, and allosteric responses. Unfortunately, the vast majority of proteins have no reliable or only partial experimentally determined 3D structures available. Such limitations may lead to presentation of a single “native” structure in visualisations instead of multiple flexible conformational variations, further perpetuating misconceptions about protein structure, folding, stability, and effects of mutations (32). The choice of protein mesh detail and representation will also impact the viewer. A low poly mesh will significantly reduce the computational burden in rendering but may compromise important scientific information, such as the specificity of a ligand binding pocket that may be essential for conveying the MoA. CG developers are however devising novel methods to visualise scientific minutiae. BioBlender is a software package that enables visual mapping of molecular surface data (charge and hydropathy) on single or multiple protein conformations that can be interpreted intuitively without the need for a legend; by combining familiar perceptual associations for these values with established methods of representing “real-world” visual or tactile experiences (33). Imperceptible field lines of electrostatic potential can be animated as curves and particle effects (flowing from positive to negative), and molecular lipophilic potentials (MLP) displayed as textures onto protein meshes (where hydrophobic regions are shiny or smooth, and hydrophilic regions are rough or dull).

Protein dynamics can range from localised movement in specific residues, to large rearrangements in domains and multiple subunits. Representing these broad spatio-temporal scales is a major challenge for biomedical animators, where protein bond vibrations and domain motions can range from femtoseconds to milliseconds respectively, and in turn many cellular processes occur in the order of seconds to minutes (34, 35). Multi-scale representations in cell landscape animations often have computer graphics performance constraints, therefore atomic resolution and motion are often sacrificed.

Cellular environments are heterogeneous, highly dynamic and densely packed. Up to 20-30% of intracellular environments can be occupied by macromolecular components (2). Molecular crowding is known to influence protein associations, diffusion, and rates of enzyme-catalysed reactions (36), yet many visualisations eschew depicting stochastic motion and extreme crowding due to fear of losing focus on the visual narrative or cognitive overload (5, 10). Furthermore, the computational expense involved with animating and rendering large numbers of meshes is a significant hurdle, however, under-representation or over-simplification has been shown to exacerbate deep rooted misconceptions, particularly amongst students (37–40). Indeed, our inherent ability to picture such numbers in real life is a challenge, and therefore it is useful to perform “sanity checks” (41).

We assessed the feasibility of replicating receptor densities based on empirical data, using the 3D computer graphics application Blender (42) and the plugin autoPACK (15). The autoPACK algorithm can fill compartmental volumes or surfaces with user defined meshes or protein meshes retrieved from the PDB, and has been previously used to generate models of HIV, blood plasma and synaptic vesicles (15, 43).

Receptor density values (number per um^2^) of 6 well-known surface biomarkers on MDA-MB-231 cells from flow cytometry data (26) were modelled on 1um^2^ area test patches. The packing simulations revealed 12650 proteins of variable sizes were easily accommodated with moderate molecular crowding (Figure 4). Surface protein diversity varies significantly between different cell types, and whilst a fairly equal distribution was modelled for this scenario, often proteins are unevenly spread or clustered in functional units. In addition, the limitations of experimental techniques ought to be scrutinised, for instance variations in antibody specificity in flow cytometry could potentially lead to under- or over-exaggeration of numbers. Often the receptor density data that is measured is actually only a snapshot in time, whereas the receptor population on a cell is highly dynamic and stochastic. Nevertheless, further exploration of experimentally derived population densities using packing algorithms such as autoPACK will facilitate understanding of surface protein distribution and organisation.

**Figure 4.**
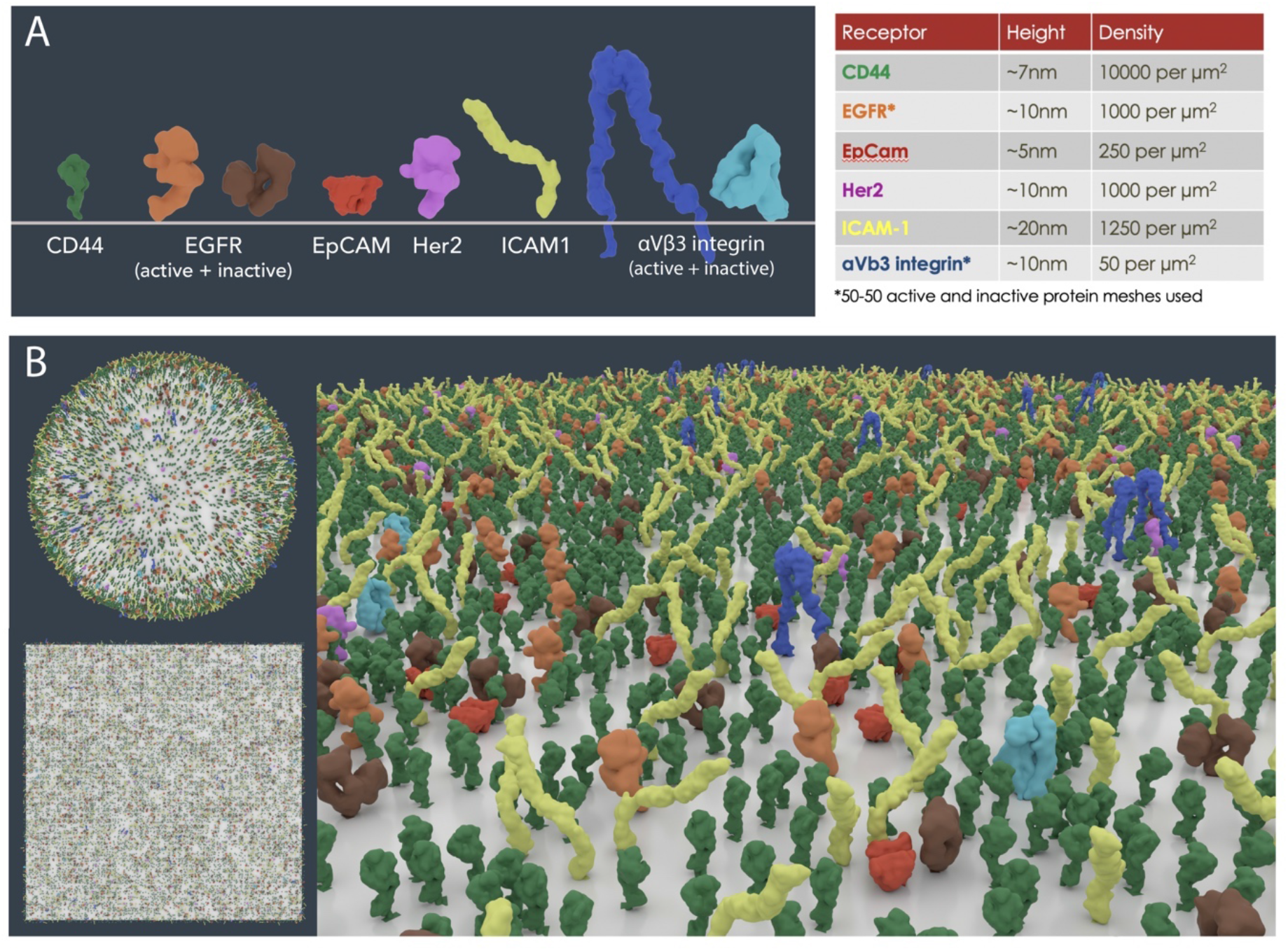
Cell surface receptor density modelling from experimental data. (A) PDB meshes of 6 well-known surface biomarkers (CD44, EGFR, EpCAM, Her2, ICAM1 and αVβ3 integrin) on MDA-MB-231 cells from flow cytometry data (26). (B) Scaled receptor meshes were distributed onto a 1μm^2^ surface area sphere and plane using the autoPACK plugin in Blender.

A negative consequence of choreographed molecular visualisations is that they can often allude to “directed intent”, the most common example being a ligand moving along undisturbed towards a receptor which subsequently leads to immediate binding. A more authentic depiction might be a random walk (constrained by kinetics and thermodynamics) through a packed environment, where the ligand undergoes many non-specific interactions before binding successfully (5, 32). Biomedical artists may be disinclined to animate unsuccessful binding events to save on animation and render time, but this will only deepen superficial understanding or promote misconceptions. Whether “visual clutter” in crowded molecular environments has a negative impact on audiences is a point of contention. Careful use of graphic devices such as titles or arrows, narration, and colour can better focus the viewer’s attention in complex visualisations and improve understanding (5, 6, 39).

### Plasma membrane lipids

All eukaryotic cells are surrounded by a plasma membrane consisting of a lipid bilayer ~4 nm thick that acts as a semi-permeable barrier between the cell interior and its environment and is a landscape where many signaling events and biological processes take place. Inside cells, specialised compartments are enclosed by lipid membranes to form discrete organelles segregating their contents from the cytoplasm (44, 45).

Lipidomics studies, predominantly through advanced mass spectrometry-based techniques and various microscopy, have identified the structure, dynamics and interactions of a diverse array of lipids within cells (46). Lipid bilayers are densely packed with approximately 5×10^6^ molecules per 1um^2^ area (47) and can encompass hundreds of different species. The most common structural lipids in eukaryotic membranes are glycerolipids (^~^65%), sterols (^~^25%) and sphingolipids (^~^10%) (31). Whilst cholesterol is generally evenly distributed between the bilayer leaflets, an asymmetric distribution of lipids contributes to their functionality (45). The inner leaflet has a greater proportion of phosphatidylethanolamine, phosphatidylserine, and phosphatidylinositol lipids, in contrast the outer leaflet is more abundant in phosphatidylcholine and sphingolipids (44).

*In vivo* imaging of membranes is experimentally difficult due to their inherent flexibility and highly dynamic fluctuations. Consequently, computational simulations mainly based on *in vitro* data have been important in understanding the heterogeneity and dynamics of plasma membranes which may exist in different phases, and the organisation of protein complexes therein. Near-atomic models of cell membranes composed of numerous lipid constitutions have been created *in silico* (46). Figure 3C depicts a typical cancer lipid bilayer with a representative complement of lipid species (31) visualised using the CHARMM-GUI Membrane Builder (48) to highlight some of the differences between the inner and outer leaflets.

Lipids are able to move freely rotationally and laterally within their leaflet and transversely between bilayers. Lateral mobility can be expressed by an experimentally determined diffusion coefficient, D. Many lipids have a D value of ^~^1 μm^2^ s^−1^ which corresponds to a lipid diffusing a distance of 2μm within 1s (49). In contrast, transverse diffusion or “flip-flop” is a far slower process that can take hours and is regulated by fippases and floppases to maintain bilayer asymmetry (49). Replicating lateral diffusion of millions of lipids at speed in whole cell and environmental molecular visualisations is not only computationally intensive, but it is questionable whether the viewer can fully observe or appreciate such minutiae in a large scene with ease and glean any real insight.

### Cellular processes

Cellular processes fall within the biological “mesoscale”, which is an intermediate scale range between nanometre-sized molecular structures and micrometre-sized cellular architecture (50, 51). An integrative modelling approach was adopted whereby information from multiple sources was combined for depiction of mesoscopic processes on the cell surface. Surface processes of interest were catagorised as either protrusions (filopodia), invaginations (caveolae, clathrin-coated pits, and macropinocytosis) and extracellular vesicles (exosomes) (Supplementary Table 2). Scanning electron microscopy (SEM) and transmission electron microscopy (TEM) data was used to calculate the approximate dimensions of typical breast cancer cells, along with sizes and densities of protrusions and invaginations (Figure 5); these measurements were in accordance with published data. Similarly, information on temporal dynamics was taken from research literature (Supplementary Table 2). Since these data fell within a broad temporal range varying from seconds to minutes, cellular features were modelled and animated using 3D software Zbrush (Pixologic) and Maya (Autodesk) (Figure 5 and Supplementary Movie 1).

**Figure 5.**
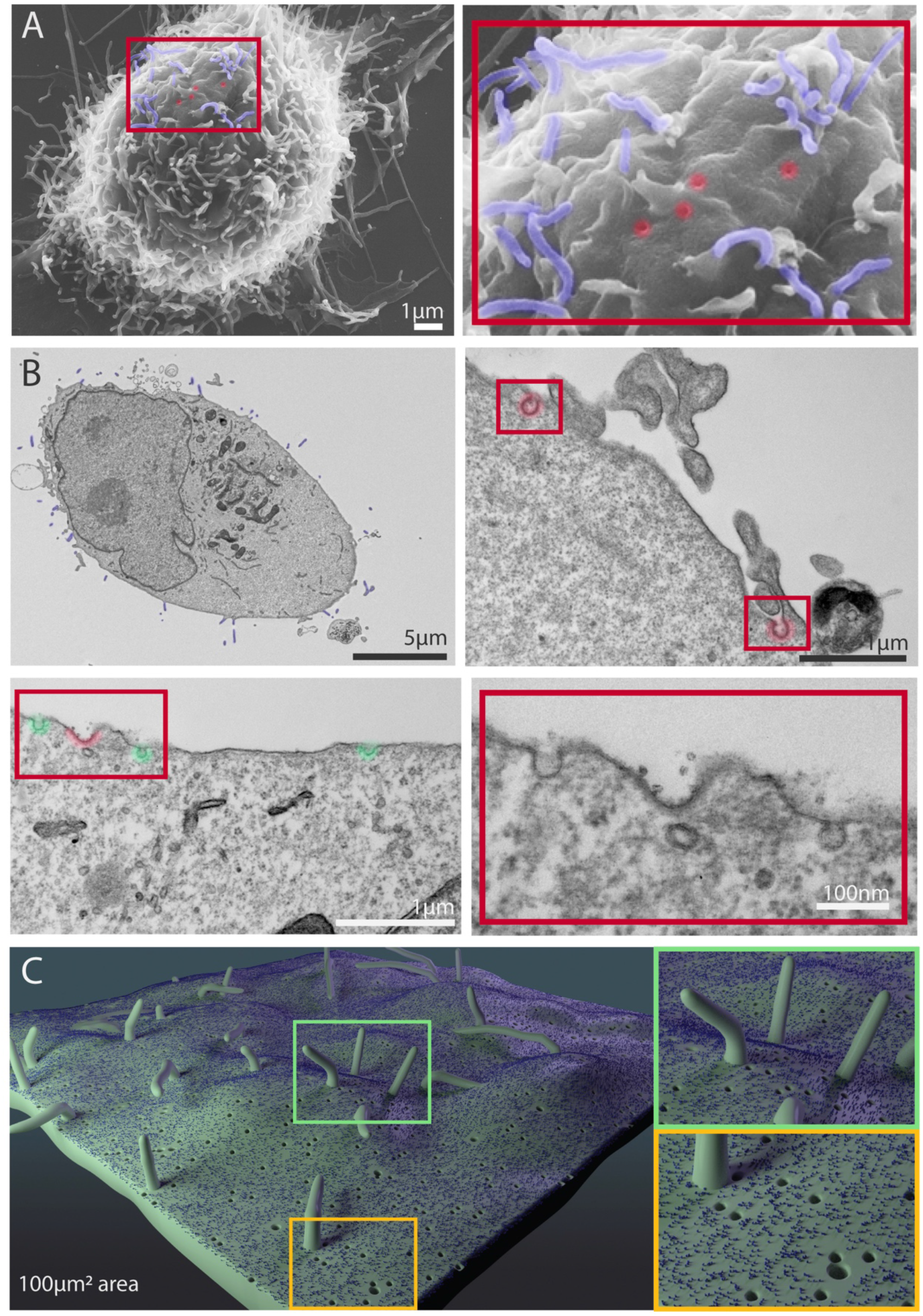
Representative SEM and TEM images of MDA-MB-231 cells showing cellular features used in Nanoscape. (A) Representative SEM images of MDA-MB-231 cells. The boxed area shows filopodia (pseudocoloured blue) and putative pits (caveolae, clathrin coated pits) in red. (B) Representative TEM images of sections of MDA-MB-231 cells. Filopodia are pseudocoloured blue, clathrin coated pits red, and caveolae green. Lower right panel shows higher magnification view of boxed area in adjacent image. (C) 3D depiction of accurate scales and densities of filopodia, caveolae, clathrin coated pits, and a representative receptor on a 100μm^2^ patch of cell membrane.

### Extracellular Matrix

The extracellular matrix (ECM) which surrounds cells *in vivo*, is an extensive complex network of structural fibers, adhesion proteins and glycosaminoglycans. Not only does it function as a scaffold for the cellular environment, but it can impact cell behaviour, processes and communication (52–54). *In vivo* the ECM consists of two biochemically and morphologically distinct forms: the basement membrane, which is a thin layer that forms between epithelial and stromal cells, and adjacent to this is the interstitial matrix, which is a more loosely organised network surrounding cells (54).

Despite its vital role, the ECM is often overlooked in many cellular and molecular visualisations and there are several reasons why. Firstly, high resolution imaging of the ECM in its native state is inherently difficult in opaque tissue. Whilst optical tissue clearing and decellularisation methods are routinely used to enhance visibility in stained tissues and organs, they are often harsh treatments which may modify the physical and chemical properties of the ECM (55). In addition, many *in vitro* models such as cells suspended in 3D gels consisting of a limited mix of collagens and matrix proteins may not always be physiologically relevant.

ECM components are often very large macromolecular complexes, and although complete protein sequences are available on Uniprot (56), many PDB structures are only of truncated or incomplete proteins. As a consequence, it is an arduous task to visualise these large ECM structures in their entirety, and some artistic license may be needed.

To visualise the ECM within a 3D breast tumour microenvironment, we collated information from the literature, including images, and sought advice from experts in the field during the conceptualisation stage (Figure 6A; Supplementary Table 2). Although there are a multitude of ECM components that make up the tumour microenvironment (54), for simplicity it was decided the focus would be only on 4 key ECM players: collagens I and IV, hyaluronic acid and fibronectin. In breast cancer, there is usually increased deposition of collagen and fibronectin, and a significant disruption of collagen IV basement membrane networks (52).

**Figure 6.**
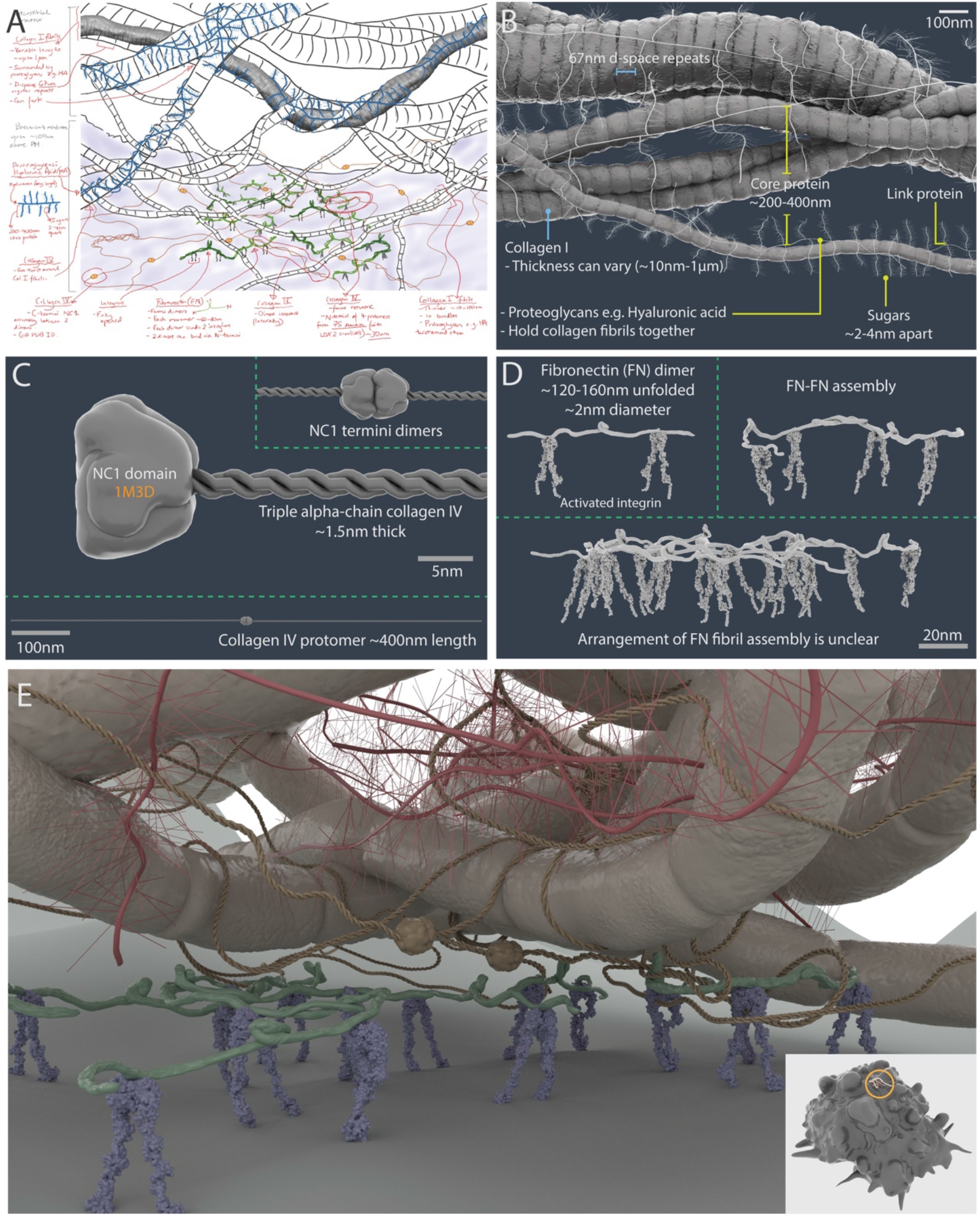
A 3D artistic impression of ECM in a tumour microenvironment. (A) Pre-production sketch layout of the scene. Scaled 3D models of collagen I fibrillar bundles and proteoglycans (e.g. hyaluronic acid) (B), collagen IV protomers and dimers (C), fibronectin dimers bound to active αVβ3 integrin (D). (E) Rendered 3D artistic interpretation of the ECM in a tumour microenvironment. Insert shows the scale of the modelled area (circle) relative to a breast cancer cell model (10μm diameter).

A stylised artistic approach was adopted to build the ECM 3D meshes using the modelling program Zbrush (Pixologic), thereby significantly reducing the polycount of large complex atomic macromolecules, which would otherwise be computationally expensive to render (Figure 6B-D and Supplementary Methods). Following advice from an ECM expert (S. Kadir personal communication with Dr Thomas Cox, Garvan Institute, July 2019), an artistic impression of a tumour microenvironment niche was built, highlighting interactions between integrin molecules on the cell surface and ECM component meshes of the basement membrane and interstitial matrix (Figure 6E).

Due to significant gaps in knowledge, many visualisation challenges were identified early on during the conceptualisation stage. Exactly how ECM proteins interact with one another to eventually form higher order structures such as fibrils, fibres and ultimately matrices, is very unclear. ECM remodelling is a highly dynamic ongoing process, which involves both proteolytic breakdown of existing matrix components by matrix metalloproteases (MMPs) and deposition of new components by cells. This intrinsic activity has significant implications on tissue development, cell migration, and pathologies including cancer (52–54). Yet there is not enough discernable data in the literature to reliably inform biomedical visualisers how to represent these processes.

Many molecular visualisations fail to show connections between the cell surface and the ECM (via cell adhesion molecules binding to ECM components) or veer away from even attempting to represent ECM density *in vivo*. This may be a combination of incomplete information about precise binding interactions, and a reluctance to complicate a scene for fear of cognitive overload. However, its frequent omission or aggressive simplification will only exacerbate a naive view of the ECM *in vivo*, whereas there is evidence that more complex molecular representations can in fact improve understanding (5, 39).

### Tumour microenvironment components

In addition to malignant cells and the ECM, a breast tumour microenvironment is made up of a complex mix of cells (including immune cells, fibroblasts, pericytes and adipocytes), blood vessels, lymphatics, and various signaling molecules (53, 57). Many studies reveal cancer cells significantly impact their environment, and interactions with non-transformed cells and the tumour vasculature have been shown to promote cancer progression (57, 58).

To build a more comprehensive tumour microenvironment, additional breast cancer cells, cancer-associated fibroblasts (CAFs) and a leaky blood vessel with an animated blood flow were incorporated into the Nanoscape scene (see Supplementary Methods and Supplementary Figure 2).

### Nanoscape user experience

Nanoscape is distinct from other published molecular and cellular visualisations, being a data-informed artistic innovation that permits user exploration and reflection of a tumour microenvironment. The Nanoscape scene was compiled in the real-time graphics engine, Unity3D and can be viewed on a desktop gaming PC (with minimum of 1080 GTX GPU). Here the user is essentially shrunk down to an equivalent height of 40nm and is able to walk on the surface of a single cancer cell within a discrete play area, surrounded by ECM, neighbouring cancer cells, CAFs, and a leaky blood vessel to observe surface proteins and cellular processes moving in real-time (Figure 7).

**Figure 7.**
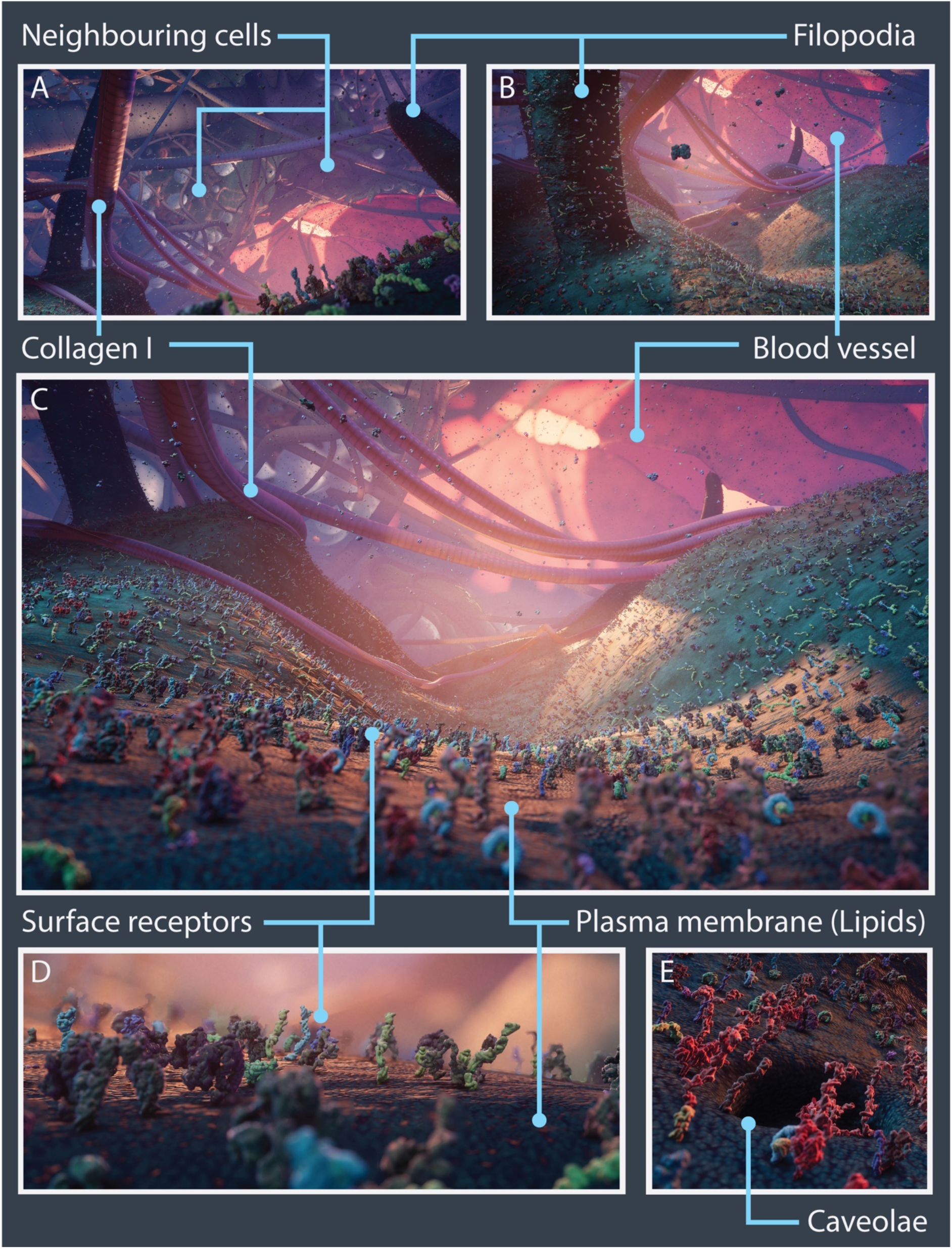
Nanoscape real-time open-world experience. Vistas from the Nanoscape real-time open-world experience with key cellular features and microenvironment components highlighted. (A – C) Panoramic views of surface receptors, the plasma membrane lipids, neighbouring cancer cells and CAFs, collagen I fibres, blood vessel and filopodia. (D) Close up of surface receptors and lipids. (E) Close up view of a surface invagination (caveolae).

Since the user experience was fundamental to the project, it became apparent that true depiction of extreme molecular crowding and the ECM had to be compromised. Extraneous complexity such as the density of soluble molecules present in the extracellular space was significantly diminished, water and ions implicit, lipid meshes were substituted with a texture to mimic their form and dynamics, and the ECM reduced to only collagen I fibres, to enable greater visibility of the landscape.

Similarly, it was computationally demanding and aesthetically unfavourable to replicate the broad temporal ranges of both molecular and cellular processes. Whist the dynamics of the major cellular processes could be animated accurately, atomic resolution was sacrificed, and protein conformational movement was appreciably slowed down to enable viewers to observe protein-protein interactions more clearly. To partially compensate for these diminutions, some artistic license was applied to evoke the “feel” of a densely packed cancerous milieu through use of colour, lighting, and sound, despite the reality that nanoscale cellular entities are below the wavelength of light and devoid of noise.

Nanoscape has significantly surpassed the level of cell surface detail and complexity compared with our previous work JTCC. Although it is not fully immersive or interactive as the VR experience of JTCC, it has been carefully choreographed to engage the viewer to appreciate the heterogeneity of a tumour microenvironment without overwhelming them with minutiae and promote understanding of some cell biology fundamentals. Future work involves development of user-defined “control features”, which empower the viewer to focus in/out of regions to switch between observing atomic detail to large cellular processes, and adjustable dials for regulating molecular density and temporal dynamics, to experience a more “authentic” cellular environment.

### Nanoscape data collection archiving

Created in 1971, the PDB remains the largest international repository for experimentally determined atomic structures. Considerable progress has been made in recent years to develop platforms and standards for archiving, validating and disseminating new biological models defined by the PDBx/mmCIF dictionary (59). Integrative or hybrid modeling structures are however currently not included in the PDB because data standards for archiving have not yet been implemented, and so the PDB-Dev was established in 2018 as a prototype archiving system (https://pdb-dev.wwpdb.org) (60). This repository contains embedded 3D viewers, links to download structures and related database entries, but also includes citations, input data and software used in the creation of models. Whilst many biomedical animators take models from such sources, clear citation of their molecular visualisation content is often limited.

The information gathered during the pre-production phase of Nanoscape is summarised in Figure 1 and Supplementary Tables 1 and 2, with references and links to PDB IDs for the proteins. Furthermore, an interactive PDF (Supplementary Figure 1) was created to document the methodology behind the creation of mMaya MoA animations for 9 receptors, associated references, and artistic approaches taken for modelling assets, along with explanatory comments. Archiving the information taken from a variety of literature, databases and communication with experts in the field, and presenting it in an accessible format will enable others to freely scrutinise or validate the work impartially, and to potentially build new, improved future versions. Selecting and curating vast amounts of information is however extremely time-consuming, requires an aptitude for interpreting scientific data, and is a constant race to keep abreast of the latest discoveries. Similarly, sustaining up-to-date versions of large-scale complex projects such as Nanoscape whenever new data becomes available or existing information is proved redundant is a huge challenge, and only reinforces the need for transparent systems of citation in scientific visualisations.

## Conclusion

Nanoscape is an innovative collaboration that has produced a multi-scale explorable 3D cellular environment. This paper sheds light on some of the technical and creative processes, decisions and sacrifices made in depicting cell surface entities and dynamics as close to experimental data as possible, whilst balancing concerns for the user experience and visual aesthetics. Although initially its main purpose was to be a unique educational and outreach tool to communicate some of the complexities of a tumour microenvironment, the final visualisation experience may in turn help experimentalists to reflect upon their own data.

Integrative modelling and visualisation of biomolecular systems and multi-scale cell models are becoming increasingly sophisticated, and educational immersive “virtual field trips” to a cell environment such as Nanoscape may provide unique insights into cell function and behaviour. As computer hardware and software continues to evolve to cope with processing enormous amounts of data, improved visual or interactive 3D representations may one day lead to novel ways for scientists to perform *in silico* experiments and potentially facilitate drug design.

An “ideal” comprehensive model would be one that fully reflects the spatio-temporal complexities and heterogeneity of an entire cell and its environment, is capable of continuous iterations and falsifiable; but to accomplish such an ambitious feat, researchers and developers from multiple scientific, computer graphics and design fields must work together.

## Supporting information

Supplementary Figure_01_Movie_01_EGFR_MoA

Supplementary Figure_01_Movie_02_Her3_MoA

Supplementary Figure_01_Movie_03_aVb3_Integrin_MoA

Supplementary Figure_01_Movie_04_VEGFR_MoA

Supplementary Figure_01_Movie_05_cKIT_MoA

Supplementary Figure_01_Movie_06_InsulinReceptor_MoA

Supplementary Figure_01_Movie_07_Tetraspanin_MoA

Supplementary Figure_01_Movie_08_TNFRSF_MoA

Supplementary Figure_01_Movie_09_GLUT1_MoA

## Acknowledgements

This work was funded by the Australian Research Council (ARC) Centre of Excellence in Convergent Bio-Nano Science and Technology (CBNS) and a National Health and Medical Research Council of Australia fellowship to RGP (APP1156489). The authors thank the Centre Director Professor Tom Davis for supporting the project, and CNBS collaborators Dr Angus Johnston (Monash University) and Professor Kristofer Thurecht (University of Queensland) for their advice. We are grateful to Dr Thomas Cox (Garvan Institute) for his expertise on the ECM and Nick Maurer for his help optimising performance out of the Unity3D game engine at the last minute. A special thanks to Mark Arrebola for advice and additional support during the project. The authors acknowledge the use of the Microscopy Australia Research Facility at the Center for Microscopy and Microanalysis at The University of Queensland and the assistance of Mr Rick Webb.

## Supplementary Data

**Supplementary Movie 1**

Animated models of cellular processes created in Maya (Autodesk): clathrin coated pits, caveolae, macropinocytosis, exosomes and filopodia. Human player model (40nm) is present for scale (also see Figure 1 for 3D models).

**Supplementary Figure 1**

Mechanism of action (MoA) animations for surface receptors: EGFR, Her3, αVβ3 integrin, VEGFR1, c-KIT, Insulin Receptor, Tetraspanin CD81 (TAPA-1), a hypothetical representative member of the TNFR super family, and GLUT1. This interactive PDF contains links to PDB structures and literary references, and embedded MoA animations. See Supplementary Methods (surface protein simulations) for information on each protein and the strategy used to create the animations.

**Supplementary Figure 2.**
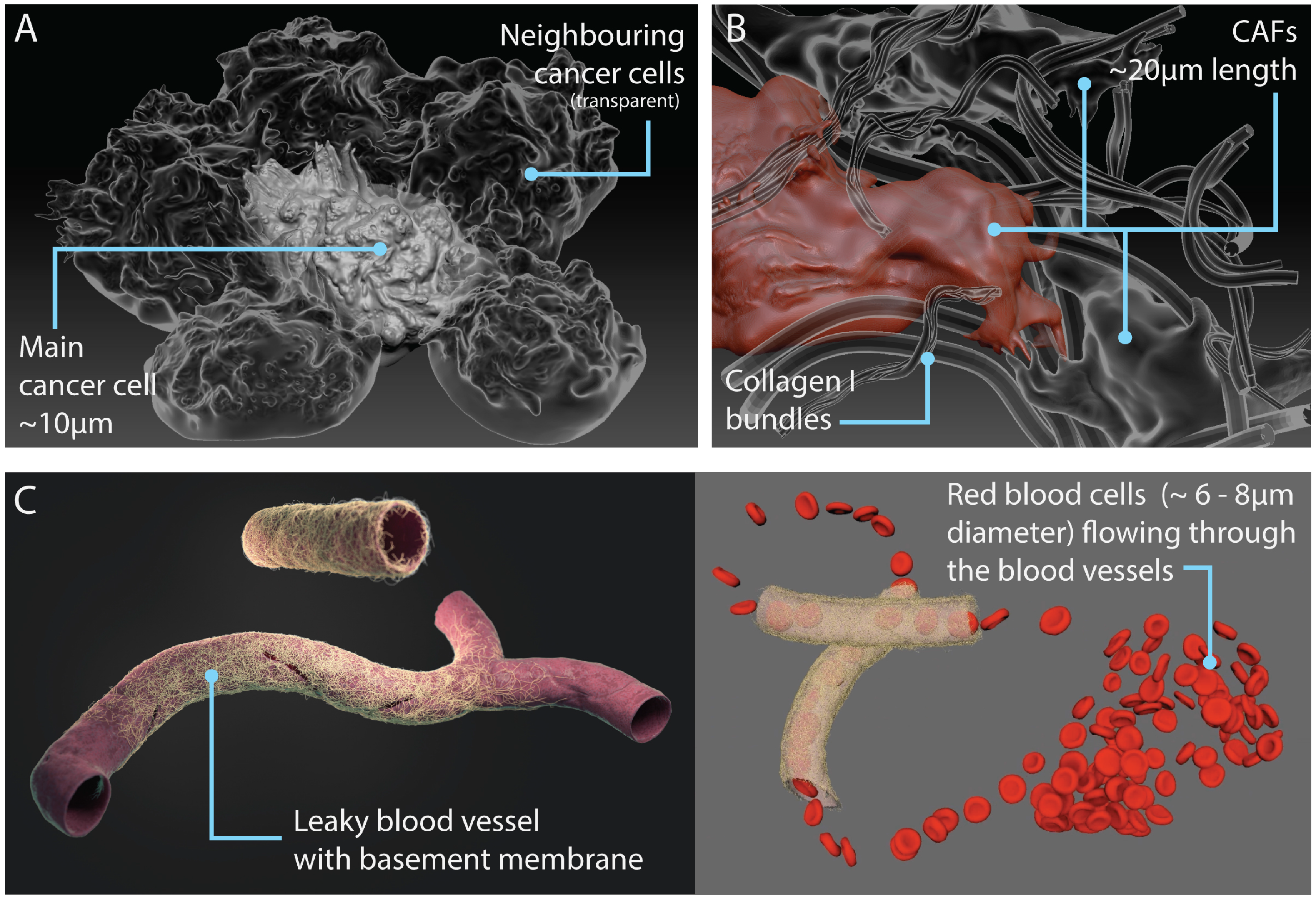
Models of components in the tumour microenvironment. (A) Additional neighbouring cancer cells (transparent) surrounding the central (play area) cancer cell. (B) Cancer-associated fibrolasts (CAFs) entangled in collagen fibres. (C) Leaky blood vessel surrounded by basement membrane mesh (left); snapshot of animation with red blood cells flowing through the vessel (right; see Supplementary Movie 2).

## Supplementary Methods

### Surface protein simulations

Mechanism of action (MoA) animations for 9 selected receptors were made using the mMaya modelling and rigging kits (30). mMaya simulations are qualitative and were used to inform the artistic design about general conformational changes and movements that may occur in receptor-ligand binding events. All rigs were “all atom no hydrogen” constructs and conformational changes were simulated by either target morphing between two end-point PDB structures (usually inactive and active states), for example EGFR (1NQL → 3NJP) (Supplementary Figure 1A and Movie 2), or manually moving handles added to selected regions of the protein rig (e.g. domains). For some proteins where only one PDB structural state was available, as with Her3 (inactive, 1M6B), movement was approximated by targeting protein domains to morph into the conformational state of another known family member (in this case active Her4 3U7U; Supplementary Figure 1A and Movie 3). In addition, simulations were inferred from MoA hypotheses published in the literature, and rig handles were applied to manipulate the movement of protein rigs to create a “hypothetical” conformation, as in the case of αVβ3 integrin extended form conformation (Supplementary Figure 1B and Movie 4). When morph targets were set between a rig and a PDB chain, the PDB chain was positioned manually to be in close proximity to the rig i.e. the c-termini of the chains were aligned as close as possible. Elastic networks were made for protein domains and the strength adjusted if necessary, to maintain the domain structures. Rig environmental dynamics were adjusted accordingly (e.g. turbulence field magnitude and damping) and simulations tested until the rig moved in a “smooth” manner. Final rig simulations were cached, and receptor surface or backbone meshes extracted. Ligand binding events were added later by keyframing the motion of the meshes manually. In some of the MoA animations, the ligand binding event was omitted and only the resultant conformational change in the protein was shown.

### ECM asset creation

Collagen I is the most abundant structural component of the interstitial membrane. Protein fibrils of varying thicknesses (10nm – 1μm) were sculpted to highlight striated 67nm d-space repeats and arranged into bundles to represent fibres. A bespoke insert mesh zbrush tool was created to wrap the proteoglycan hyaluronic acid around collagen I structures (Figure 6B and E).

Similarly, an insert mesh zbrush tool was made for modelling Collagen IV protomers, which consisted of three intertwining α-chains that form a triple helix 400nm in length. Two collagen IV protomers were joined head-to-head via NC1 dimers (PDB structure 1M3D) at the C-termini, and four collagen N-termini overlaid to form the 7S domain (28nm overlaps), to create an extensive branched mesh network (Figure 6C and E).

Fibronectin monomers are made up of three repeating units (FN types I, II, III) and usually form dimers linked by a pair of disulfide bonds at their c-termini (61). Fibronectin dimers are long and can form fibrils (ranging from ~133-190 nm) (62) however the precise molecular arrangement and their associations with multiple binding partners is still unclear. Alternative splicing can lead to over 20 protein variants in humans (61); therefore, a simplified dimer mesh was modelled which was shown only bound to integrin (Figure 6D and E).

### Cancer cells, CAFs and blood vessel

Tumour microenvironment components: additional neighbouring cancer cells (~10μm diameter), cancer-associated fibroblasts (CAFs) (~20μm length) and a leaky blood vessel (~10μm diameter) with red blood cells (~6-8μm diameter) were modelled in Zbrush based on various microscopy images taken from the literature (See supplementary Figure 2). The blood flow in the vessel was animated in Maya (Supplementary Movie 2).

---

**Supplementary Table 1.** Breast cancer-associated surface protein structures from the RSCB protein data bank (PDB), their PDB ID, associated mechanism of action references, and supplementary movies for selected proteins are indicated with an asterix (see Figure 2 and Supplementary Figure 1 for details).

| Receptor/soluble protein | Ligand | PDB Structures | Description |
| --- | --- | --- | --- |
| EGFR (HER1, ERBB1)* | EGF or TGF-α | 1NQL | EGFR monomer (unliganded) |
|  |  | 3NJP | EGFR dimer (EGF bound) |
| | | 1MOX | EGFR (TGF-α bound) |
| HER2 (ERBB2) | No known ligand | 1S78 | Her2 Monomer (unliganded) |
| HER3 (ERBB3)* | Neuregulins (NRG1 + NRG2) | 1M6B | Her3 Monomer (unliganded) |
| HER4 (ERBB4) | NRG1 | 2AHX | Her4 Monomer (unliganded) |
|  |  | 3U7U | Her4 Monomer (NRG1 bound) |
| Integrin αVβ3* | Various e.g. ECM proteins | 4G1M | alpha V beta 3 structure (inactive) |
| VEGFR-1* | VEGF-A | VEGFR-1 D1-7 VEGF-A composite model | Full-length structure of the VEGFR-1 ectodomain (VEGF bound) |
| c-KIT* | SCF | 2EC8 | c-KIT monomer (unliganded) |
|  |  | 2E9W | c-KIT homodimer (SCF bound) |
| Insulin Receptor (IR)* | Insulin | 4ZXB | Insulin Receptor heterodimer (unliganded) |
|  |  | 6CEB | Insulin Receptor heterodimer ectodomain (two insulin molecules bound) |
| Insulin-like growth factor receptor (IGF1R) | IGF-I | 5U8Q | Type 1 insulin-like growth factor receptor heterodimer (IGF-I bound) |
| MHC I (aka HLA-B27) | N/A (presents antigens) | 1HSA | HLA-B27 and beta-2 microglobulin (model peptide sequence bound) |
| TLR4-MD2 complex | LPS (lipopolysaccharide) | 3FXI | Toll-like receptor 4 and MD-2 (lipopolysaccharide of Gram-negative bacteria bound) |
| TLR1-TLR2 heterodimer | Lipopeptide | 2Z7X | TLR1-TLR2 heterodimer (tri-acylated lipopeptide bound) |
| CXCR4 | CXCL12 (SDF-1) | 3OE0 | CXCR4 chemokine receptor in complex with small molecule antagonist IT1t |
|  |  | 2K05 | SDF1 in complex with the CXCR4 N-terminus |
| tetraspanin CD81* (TAPA-1) | Cholesterol | 5TCX | Tetraspanin CD81 |
| TNFR1* | TNF, LTα, LTβ | 4MSV | FASL and DcR3 complex (used instead as TNFR1 is incomplete) |
|  |  | 2NA7 | Transmembrane domain of human Fas/CD95 death receptor |
|  |  | Stem Linker | Constructed in mMaya from sequence |
| LDL receptor-PCSK9 complex | lipoproteins | 1N7D | Extracellular domain of the LDL receptor at endosomal pH (no lipoprotein bound) |
|  |  | 3P5C | PCSK9-LDLR structure at neutral pH (no lipoprotein bound) |
| CSF1R | CSF1 | 4WRL | Dimeric CSF-1 bound to two CSF-1R molecules (D1–D3 part only) |
|  |  | 4WRM | CSF-1R monomer (CSF-1 bound) |

Supplementary Table 1
|  |  |  |  |
| --- | --- | --- | --- |
| Transferrin Receptor (TfR) | Transferrin (Tf) | 1SUV | Transferrin Receptor (Serotransferrin, N-lobe and C-lobes bound) |
| EphA2 | EphrinA5 | 2X10 | EphA2 ectodomain (monomer) |
|  |  | 3MX0 | EphA2 ectodomain (in complex with ephrin A5) |
| EphA4 | EphrinA5 (and others) | 4M4P | EphA4 ectodomain (monomer) |
|  |  | 4M4R | Epha4 ectodomain (complex with ephrin A5) |
| MMP-2 (Gelatinase A) | Various e.g. ECM proteins | 1CK7 | MMP-2 (Gelatinase A) |
| MMP-9 (Gelatinase B) | Various e.g. ECM proteins | 1L6J | MMP-9 (Gelatinase B) |
| TIMP-2 (tissue inhibitors of metalloproteinases-2) | MMPs | 1GXD | proMMP-2/TIMP-2 complex |
| CD44 | glycosaminoglycan hyaluronan, collagens, osteopontin, MMPs | 2JCP | Hyaluronan binding domain of murine CD44 |
|  |  | Standard stem region | Constructed in mMaya from sequence |
| EpCAM | EpCAM (on other cells) | 4MZV | EpCAM cis-dimer |
| N-Cadherin | N-Cadherin (on other cells) | 3Q2W | N-Cadherin trans dimer |
| E-Cadherin | E-Cadherin (on other cells) | 3Q2V | E-Cadherin trans dimer |
| ICAM | LFA-1 (an integrin present on leukocytes) | 1IC1 | ICAM-1 D1-D2 domains |
|  |  | 1P53 | ICAM-1 D3-D5 domains |
| Glucose transporter GLUT-1* | Glucose | 4PYP | GLUT1 (inward-open; glucose-bound) |
|  |  | 4ZWC | GLUT3 (outward-open; maltose-bound) |
| Potassium Channel Kv1.2 | K+ | 3LUT | Full-length Shaker Potassium Channel Kv1.2 |
| Na+/K+ pump | Na+, K+ | 2ZXE | Sodium-Potassium pump |
| ABCA1 | phospholipids and cholesterol | 5XJY | Lipid Exporter ABCA1 |

\* Supplementary MoA animation movies are available (see Supplementary Figure 1 for further details)

**Supplementary Table 2.** Key features of cellular structures and processes in Nanoscape with examples detailing properties such as dimensions, densities, temporal dynamics, and their associated literature references (also see Figure 1 for 3D models).

|  | Feature | Examples | Dimensions | Density | Temporal dynamics |
| --- | --- | --- | --- | --- | --- |
| Structures | Membrane bound receptors<sup>1</sup> | EGFR, Integrins, VEGFR | ~ 5-20nm length;<br>~ 1-5nm diameter | Protein specific<br>~ 10-10<sup>4</sup> per $\mu\text{m}^2$ | Protein specific<br><sup>†</sup>D ~ 10<sup>-3</sup>-1$\mu\text{m}^2$/s;<br>Transitions between protein states ~ 1-100$\mu\text{s}$ |
|  | Soluble proteins | EGF, TNF, MMPs |  |  |  |
| | Membrane transport proteins<sup>2</sup> | GLUT4, K<sup>+</sup> channel, Na<sup>+</sup>/K<sup>+</sup> pump | ~ 4-10nm length | ~ 10<sup>4</sup> per $\mu\text{m}^2$ | ~ 10<sup>0</sup>-10<sup>7</sup>/s transport rate |
|  | Extracellular matrix<sup>3</sup> | Collagens, Fibronectin, Hyaluronic acid | Variable: from large fibres to smaller glycoproteins | Variable | Very low mobility relative to proteins |
| | Plasma membrane<sup>4</sup> | Phospholipids | ~ 2nm length; ~ 0.25-0.5nm<sup>2</sup> cross-sectional area | ~ 5 x 10<sup>6</sup> per $\mu\text{m}^2$ | <sup>†</sup>D ~ 1$\mu\text{m}^2$/s |
| Processes | Protrusions<sup>5</sup> | Filopodia | ~ 1-5$\mu\text{m}$ length;<br>~ 150-200nm diameter | ~ 0.3 per $\mu\text{m}^2$ | ~ 25-50nm/s protrusion rate |
| | Endocytosis | Caveolae<sup>6</sup> | ~ 65nm mean diameter;<br>~ 0.0067$\mu\text{m}^2$ area | ~ 10 per $\mu\text{m}^2$ | ~ 30s to minutes |
| | | Clathrin mediated endocytosis<sup>7</sup> | ~ 110nm mean diameter;<br>~ 0.0190$\mu\text{m}^2$ area | ~ 0.8 per $\mu\text{m}^2$ | ~ 30-60s |
| | | Macropinocytosis<sup>8</sup> | ~ 0.2-5$\mu\text{m}$ diameter | ? | ~ 120s |
|  | Extracellular vesicles | Exosomes<sup>9</sup> | ~ 40-150nm diameter | ? | ? |

<sup>†</sup>D = Diffusion coefficient (microscopically determined by the velocity of the molecule and the mean time between collisions)

## References

**1. Membrane bound receptors and soluble proteins:**

Milo, R., Jorgensen, P., Moran, U. et al. BioNumbers--the database of key numbers in molecular and cell biology. Nucleic acids research, 38 (Database issue), D750-D753; (2010). Available from: https://doi.org/10.1093/nar/gkp889

**2. Membrane transport proteins:**

Milo, R., Jorgensen, P., Moran, U. et al. BioNumbers--the database of key numbers in molecular and cell biology. Nucleic acids research, 38 (Database issue), D750-D753; (2010). Available from: https://doi.org/10.1093/nar/gkp889. Chapter IV: Rates and Duration; http://book.bionumbers.org/how-many-ions-pass-through-an-ion-channel-per-second/

Itzhak, D., Tyanova, S., Cox, J. et al. Global, quantitative and dynamic mapping of protein subcellular localization. Elife, 5, pii: e16950; p.9 top paragraph, (2016). Available from: https://doi.org/10.7554/eLife.16950.

Gennis, R. Biomembranes: Molecular Structure and Function. Springer-Verlag, N.Y., p.274 table 8.3, (1989). Available from: https://www.springer.com/gp/book/9781475720679

**3. Extracellular Matrix:**

Frantz, C., Stewart, K. & Weaver, V. The extracellular matrix at a glance. Journal of Cell Science, 123(24): p. 4195, (2010). Available from: https://doi.org/10.1242/jcs.023820

Insua-Rodríguez, J. & Oskarsson, T. The extracellular matrix in breast cancer. Advanced Drug Delivery Reviews, 97: p. 41-55, (2016). Available from: https://doi.org/10.1016/j.addr.2015.12.017

Mouw, J., Ou, G. & Weaver, V. Extracellular matrix assembly: a multiscale deconstruction. Nature Reviews Molecular Cell Biology, 15(12): p. 771-785, (2014). Available from: https://doi.org/10.1038/nrm3902

Pankov, R. & Yamada, K. Fibronectin at a glance. Journal of Cell Science, 115(20): p. 3861, (2002). Available from: https://doi.org/10.1242/jcs.00059

Früh, S., Schoen, I., Ries, J. et al. Molecular architecture of native fibronectin fibrils. Nature Communications, 6, 7275, (2015). Available from: https://doi.org/10.1038/ncomms8275

**4. Plasma membrane lipids:**

Milo, R., Jorgensen, P., Moran, U. et al. BioNumbers--the database of key numbers in molecular and cell biology. Nucleic acids research, 38 (Database issue), D750-D753; (2010). Available from: https://doi.org/10.1093/nar/gkp889. Chapter II: Concentrations and Absolute Numbers; http://book.bionumbers.org/what-lipids-are-most-abundant-in-membranes/

Alberts, B., Johnson, A., Lewis, J. et al. Molecular Biology of the Cell. 4th edition. New York: Garland Science; (2002). The Lipid Bilayer. Available from: https://www.ncbi.nlm.nih.gov/books/NBK26871/

Rawicz, W., Olbrich, K., McIntosh, T. et al. Effect of chain length and unsaturation on elasticity of lipid bilayers. Biophysical Journal, 79(1):328-39. p.332 table 1, (2000). Available from: https://doi.org/10.1016/S0006-3495(00)76295-3

Brügger, B., Glass, B., Haberkant, P. et al. The HIV lipidome: a raft with an unusual composition. Proceedings of the National Academy of Sciences of the United States of America, 103(8):2641-6. p.2644 right column 2nd paragraph, (2006). Available from: https://doi.org/10.1073/pnas.0511136103

**5. Filopodia density and dimensions:**

Measured from scanning electron micrographs, see Figure 5.

Mallavarapu, A. & Mitchison, T. Regulated Actin Cytoskeleton Assembly at Filopodium Tips Controls Their Extension and Retraction. The Journal of Cell Biology, 146 (5) 1097-1106; (1999). Available from: https://doi.org/10.1083/jcb.146.5.1097

**6. Caveolae density, dimensions and temporal dynamics:**

Parton, R., McMahon, K. & Wu Y. Caveolae: Formation, dynamics, and function. Current Opinion in Cell Biology, 65:8-16, (2020). Available from: https://doi.org/10.1016/j.ceb.2020.02.001

Parton, R. Ultrastructural localization of gangliosides; GM1 is concentrated in caveolae. Journal of Histochemistry & Cytochemistry, 42(2):155-166, (1994). Available from: https://doi.org/10.1177/42.2.8288861

Richter, T., Floetenmeyer, M., Ferguson, C. et al. High-resolution 3D quantitative analysis of caveolar ultrastructure and caveola-cytoskeleton interactions. Traffic, 9(6):893-909, (2008). Available from: https://doi.org/10.1111/j.1600-0854.2008.00733.x

Parton, R., Kozlov, M. & Ariotti, N. Caveolae and lipid sorting: Shaping the cellular response to stress. Journal of Cell Biology, 219(4):e201905071, (2020). Available from: https://doi.org/10.1083/jcb.201905071

Boucrot, E., Howes, M., Kirchhausen, T. et al. Redistribution of caveolae during mitosis. Journal of Cell Science, 124(Pt 12):1965-1972, (2011). Available from: https://doi.org/10.1242/jcs.076570

Pelkmans, L. & Zerial, M. Kinase-regulated quantal assemblies and kiss-and-run recycling of caveolae. Nature, 436(7047):128-133, (2005) Available from: https://doi.org/10.1038/nature03866

**7. Clathrin mediated endocytosis density, dimensions and temporal dynamics:**

Parton, R. Ultrastructural localization of gangliosides; GM1 is concentrated in caveolae. Journal of Histochemistry & Cytochemistry, 42(2):155-166, (1994). Available from: https://doi.org/10.1177/42.2.8288861

Edeling, M., Smith, C. & Owen, D. Life of a clathrin coat: insights from clathrin and AP structures. Nature Reviews Molecular Cell Biology, 7(1):32-44, (2006). Available from: https://doi.org/10.1038/nrm1786

Doherty, G. & McMahon, H. Mechanisms of endocytosis. Annual Review of Biochemistry, 78:857-902, (2009). Available from: https://doi.org/10.1146/annurev.biochem.78.081307.110540

McMahon, H. & Boucrot, E. Molecular mechanism and physiological functions of clathrin-mediated endocytosis. Nature Reviews Molecular Cell Biology, 12(8):517-533, (2011). Available from: https://doi.org/10.1038/nrm3151

Merrifield, C., Feldman, M., Wan, L. et al. Imaging actin and dynamin recruitment during invagination of single clathrin-coated pits. Nature Cell Biology, 4(9):691-698, (2002). Available from: https://doi.org/10.1038/ncb837

Saffarian, S. & Kirchhausen, T. Differential evanescence nanometry: live-cell fluorescence measurements with 10-nm axial resolution on the plasma membrane. Biophysical Journal, 94(6):2333-2342, (2008). Available from: https://doi.org/10.1529/biophysj.107.117234

Kirchhausen, T. Imaging endocytic clathrin structures in living cells. Trends in cell biology, 19,11: 596-605, (2009). Available from: https://doi.org/10.1016/j.tcb.2009.09.002

Taylor, M., Perrais, D. & Merrifield, C. A high precision survey of the molecular dynamics of mammalian clathrin-mediated endocytosis. PLoS Biology, 9(3):e1000604, (2011). Available from: https://doi.org/10.1371/journal.pbio.1000604

Cocucci, E., Aguet, F., Boulant, S. et al. The first five seconds in the life of a clathrin-coated pit. Cell, 150(3):495-507, (2012). Available from: https://doi.org/10.1016/j.cell.2012.05.047

**8. Macropinocytosis dimensions:**

Lim, J. & Gleeson, P. Macropinocytosis: an endocytic pathway for internalising large gulps. Immunology & Cell Biology, 89: 836-843, (2011). Available from: https://doi.org/10.1038/icb.2011.20

Condon, N., Heddleston, J., Chew, T. et al. Macropinosome formation by tent pole ruffling in macrophages. Journal of Cell Biology, 217(11):3873-3885, (2018). Available from: https://doi.org/10.1083/jcb.201804137

**9. Exosomes dimensions:**

Skotland, T., Sandvig, K. & Llorente, A. Lipids in exosomes: Current knowledge and the way forward. Progress in Lipid Research, 66:30-41, (2017). Available from: https://doi.org/10.1016/j.plipres.2017.03.001

## Supplementary Figure 1

Mechanism of action animations for 9 selected surface receptors. See Supplementary Methods for details. Links for PDBs, reference papers and MoA animations are embedded.

- Her Family (EGFR and Her3)
- αVβ3 integrin
- VEGFR1
- c-KIT
- Insulin Receptor
- Tetraspanin CD81 (TAPA-1)
- TNFR super family
- GLUT1

N.B. For this initial submission the MoA animations have not been embedded in Supplementary Figure 1 as it exceeds the file size requirements but are available as mp4 files for viewing.

### Her Family

- All Her family members have 4 extracellular domains (I - IV).
- In the monomeric tethered conformation (EGFR, Her3 and Her4), the dimerization arm (in domain II) is completely occluded by intramolecular interactions with domain IV.
- Ligand binding causes a conformational change ~ 130° rotation of domains I + II with respect to domains III + IV, and leads to dimerisation (homo- and heterodimerisation with Her family members).
- The dimerization arm of one monomer interacts with the corresponding element of the dimer partner.
- Her2 exists in the extended conformation without ligand binding.

#### EGFR (Her1, ERBB1)

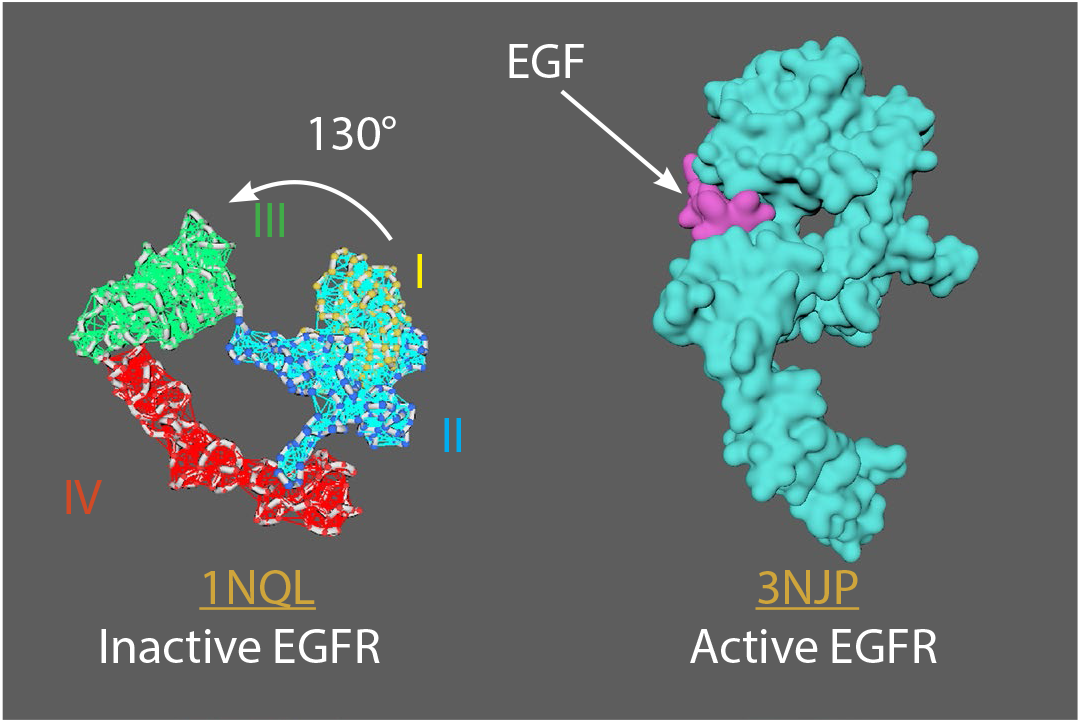

**EGFR MoA animation strategy:**

1NQL^1^ = inactive EGFR (monomer)

3NJP^2^ = active EGFR (dimer)

Rigged 1NQL with mMaya rigging kit, created elastic networks for each domain: I+II combined, III, and IV. Set 3NJP as a target for 1NQL to morph into, and cached the animation.

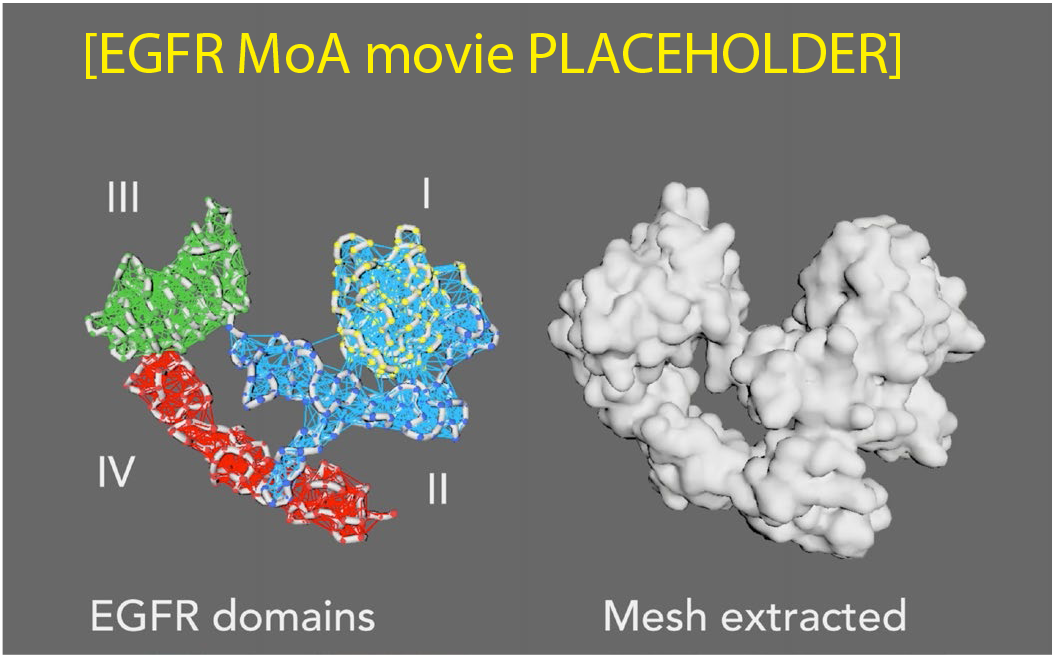

#### Her3 (ERBB3)

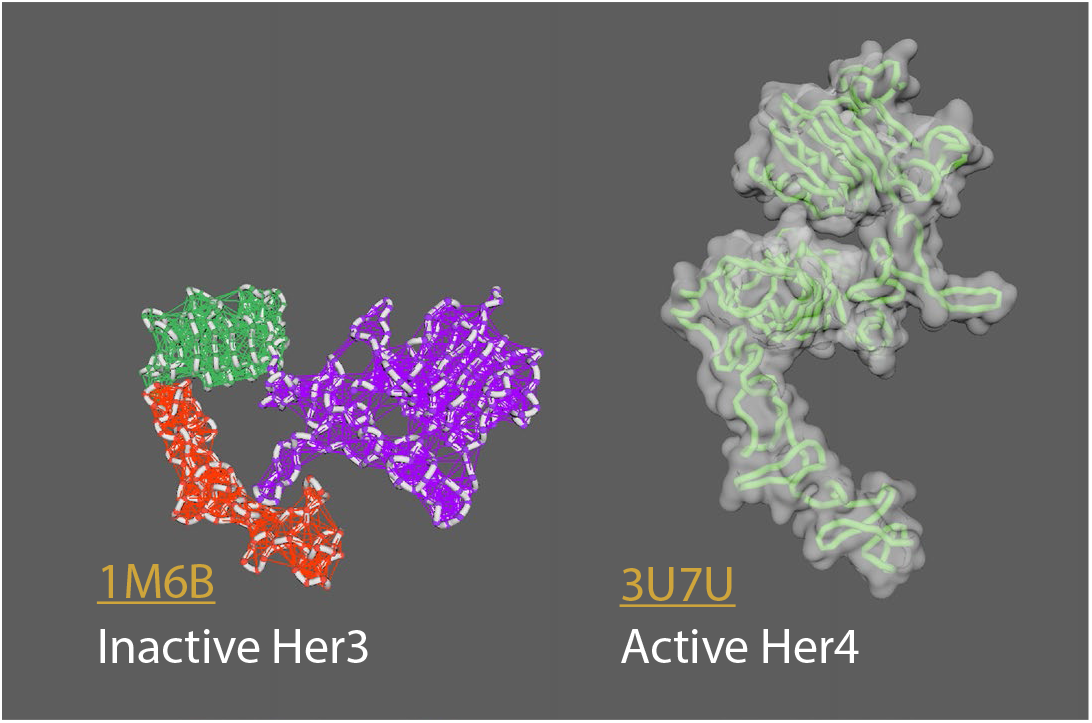

**Her3 MoA animation strategy:**

1M6B^3^ = inactive Her3 (monomer)

3U7U^4^ = active Her4 (dimer)

Rigged 1M6B with mMaya rigging kit, created elastic networks for each domain: I+II combined, III, and IV. Set 3U7U as a target for 1M6B to morph into, and cached the animation.

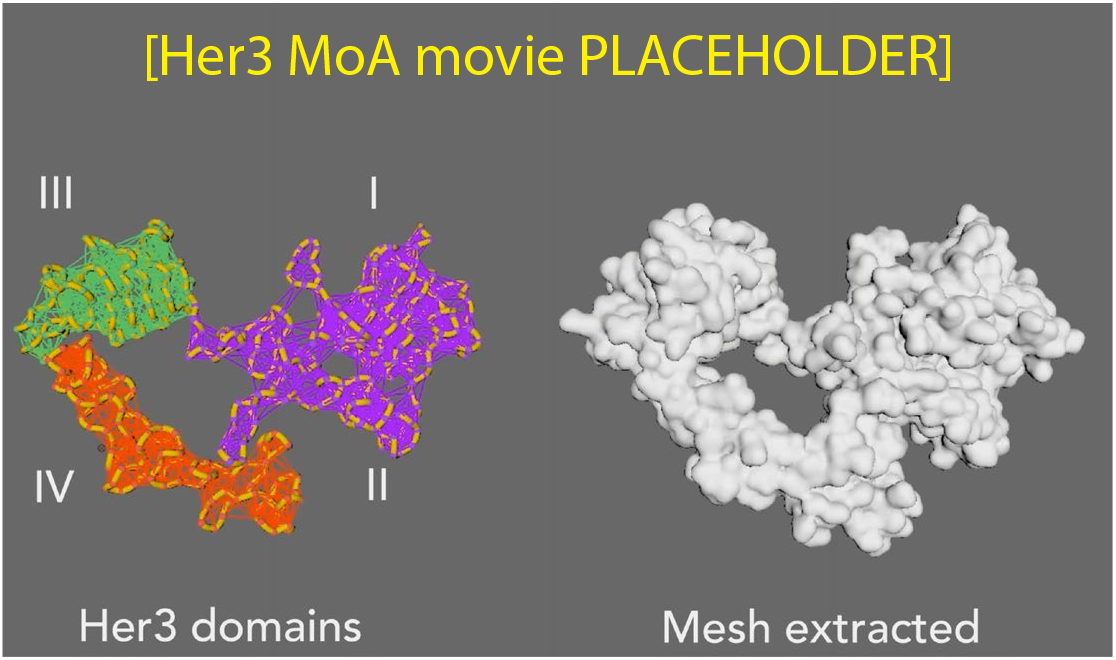

##### References

1. Ferguson, K., Berger, M., Mendrola, J. et al. EGF Activates Its Receptor by Removing Interactions that Autoinhibit Ectodomain
Dimerization. Molecular Cell, 11 (2), 507-517. (2003). https://doi.org/10.1016/S1097-2765(03)00047-9

2. Lu, C., Mi, L., Grey, M. et al. Structural Evidence for Loose Linkage between Ligand Binding and Kinase Activation in the Epidermal Growth Factor Receptor. Molecular and Cellular Biology, 30 (22) 5432-5443 (2010); https://doi.org/10.1128/MCB.00742-10

3. Cho, H. and Leahy, D. Structure of the Extracellular Region of HER3 Reveals an Interdomain Tether. Science, 297 (5585), 1330, (2002). https://doi.org/10.1126/science.1074611

4. Liu, P., Cleveland IV, T., Bouyain, S. et al. A single ligand is sufficient to activate EGFR dimers. Proceedings of the National Academy of Sciences, 109 (27), 10861, (2012). https://doi.org/10.1073/pnas.1201114109

5. Kovacs, E., Zorn, J., Huang, Y. et al. A structural perspective on the regulation of the epidermal growth factor receptor. Annual review of biochemistry, 84, 739–764, (2015). https://doi.org/10.1146/annurev-biochem-060614-034402

### αVβ3 integrin

- Three major integrin conformational states: (A) bent with closed headpiece, (B) extended with a closed headpiece, and (C) extended with an open headpiece.
- Switchblade model: conformational change from bent to extended
- Extracellular domains: 4 in alpha; 8 in beta chains
- Bent conformation at knees/genua: between thigh and calf-1 in alpha5; between EGF1 and EGF2 in beta3
- Ligands vary (e.g. ECM proteins)

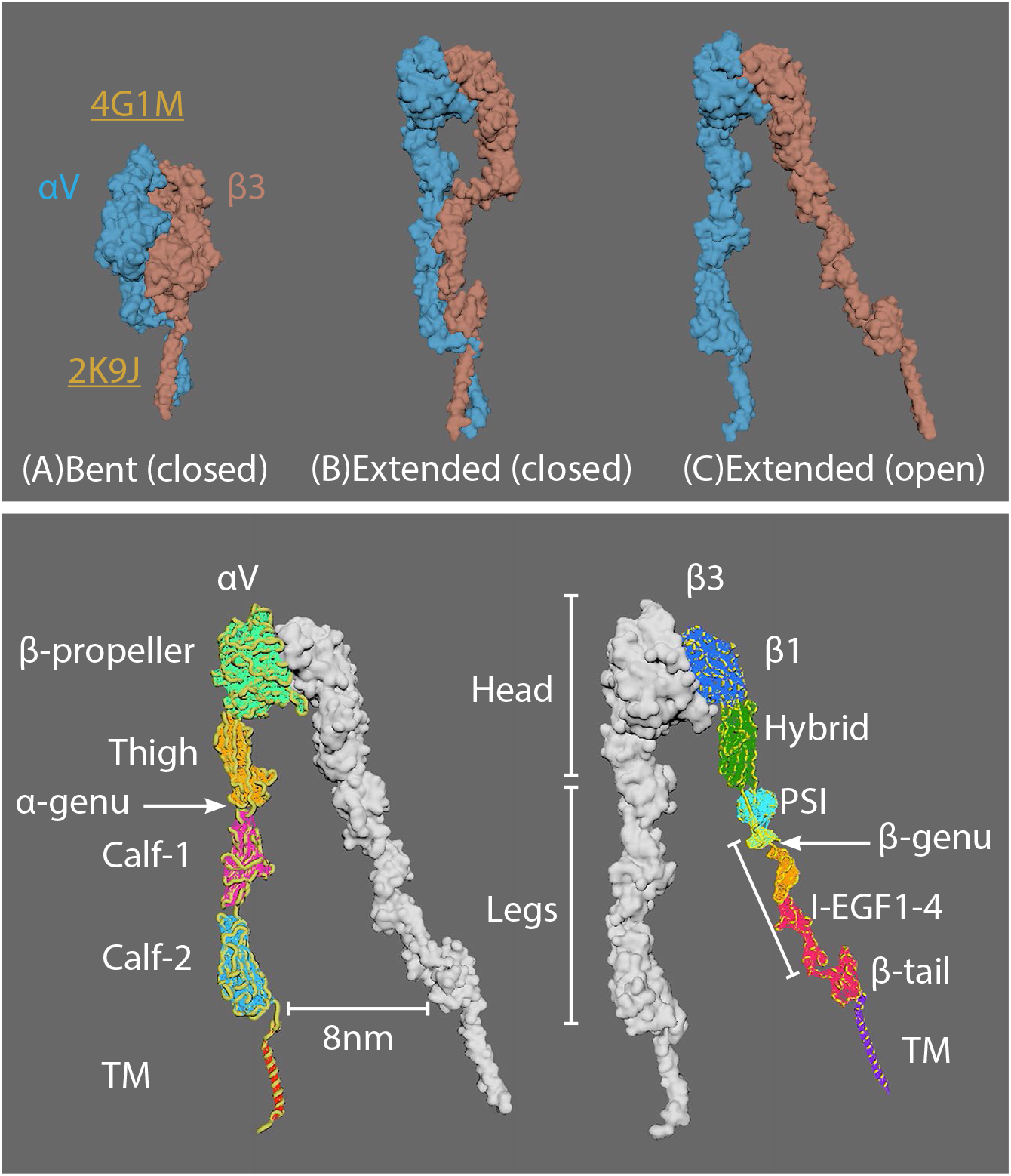

**MoA animation strategy:**

4G1M^1^ = αVβ3 inactive bent conformation

2K9J2 = transmembrane domain of integrin αIIb-β3

Connected 2K9J to the C-term of αVβ3 integrin (4G1M) using the mMaya modelling kit. New PDB created was rigged and elastic networks made for all extracellular domains. As no Extended structures are available artistic licence was used to create hypothetical structures. Handles were created for the Head portion of αV and β3 to manually open the mesh into the extended closed conformation, and a second handle created for the I-EGFR1-4 and β-tail to move ~8nm into the extended open conformation.

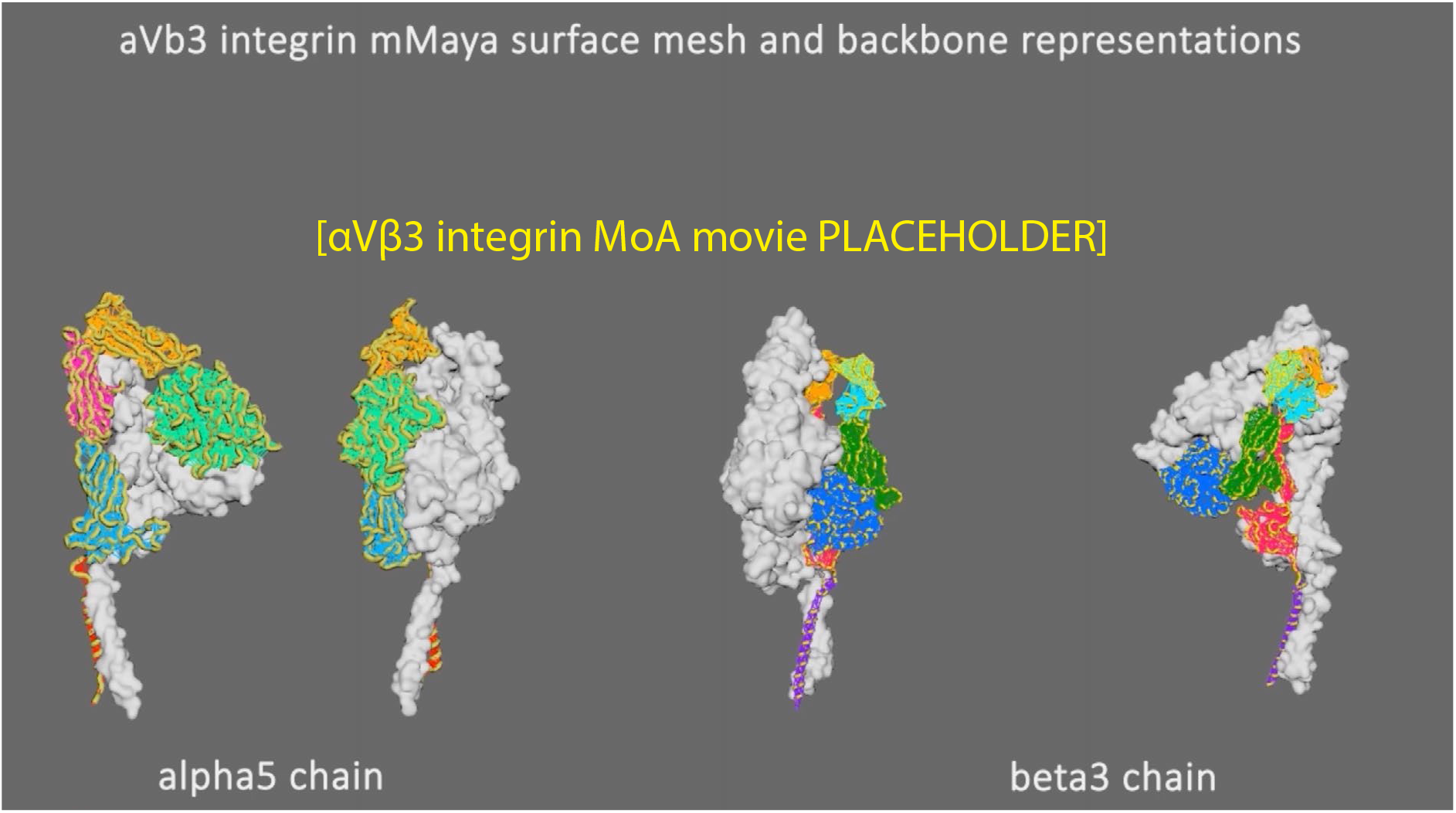

#### References

1. Dong, X., Mi, L., Zhu, J. et al. αVβ3 Integrin Crystal Structures and Their Functional Implications. Biochemistry, 51 (44), 8814-8828 (2012). https://doi.org/10.1021/bi300734n

2. Lau, T., Kim, C., Ginsberg, M. et al. The structure of the integrin αIIbβ3 transmembrane complex explains integrin transmembrane signalling. The EMBO Journal, 28: 1351-1361, (2009). https://doi.org/10.1038/emboj.2009.63

3. Zhu, J., Luo B., Xiao, T. et al. Structure of a Complete Integrin Ectodomain in a Physiologic Resting State and Activation and Deactivation by Applied Forces. Molecular Cell, 32 (6), 849-861 (2008). https://doi.org/10.1016/j.molcel.2008.11.018

4. Chen, W., Lou, J., Hsin, J. et al. Molecular Dynamics Simulations of Forced Unbending of Integrin αVβ3. PLOS Computational Biology 7(2): e1001086, (2011). https://doi.org/10.1371/journal.pcbi.1001086

### VEGFR1

- VEGFR1 has 7 Ig-like domains (D1-D7) and binds VEGF-A (dimer)
- Full composite dimer (available from Markovic-Mueller et al. 2017) built from PDBs 5T89 (VEGFR-1 Domains 1-6) and 3KVQ (VEGFR-2 Domain 7)
- D2 and D3 ectodomains are binding sites for VEGF-A
- D3/VEGF-A binding triggers interactions between D4–5 and D7 in VEGFR homodimers

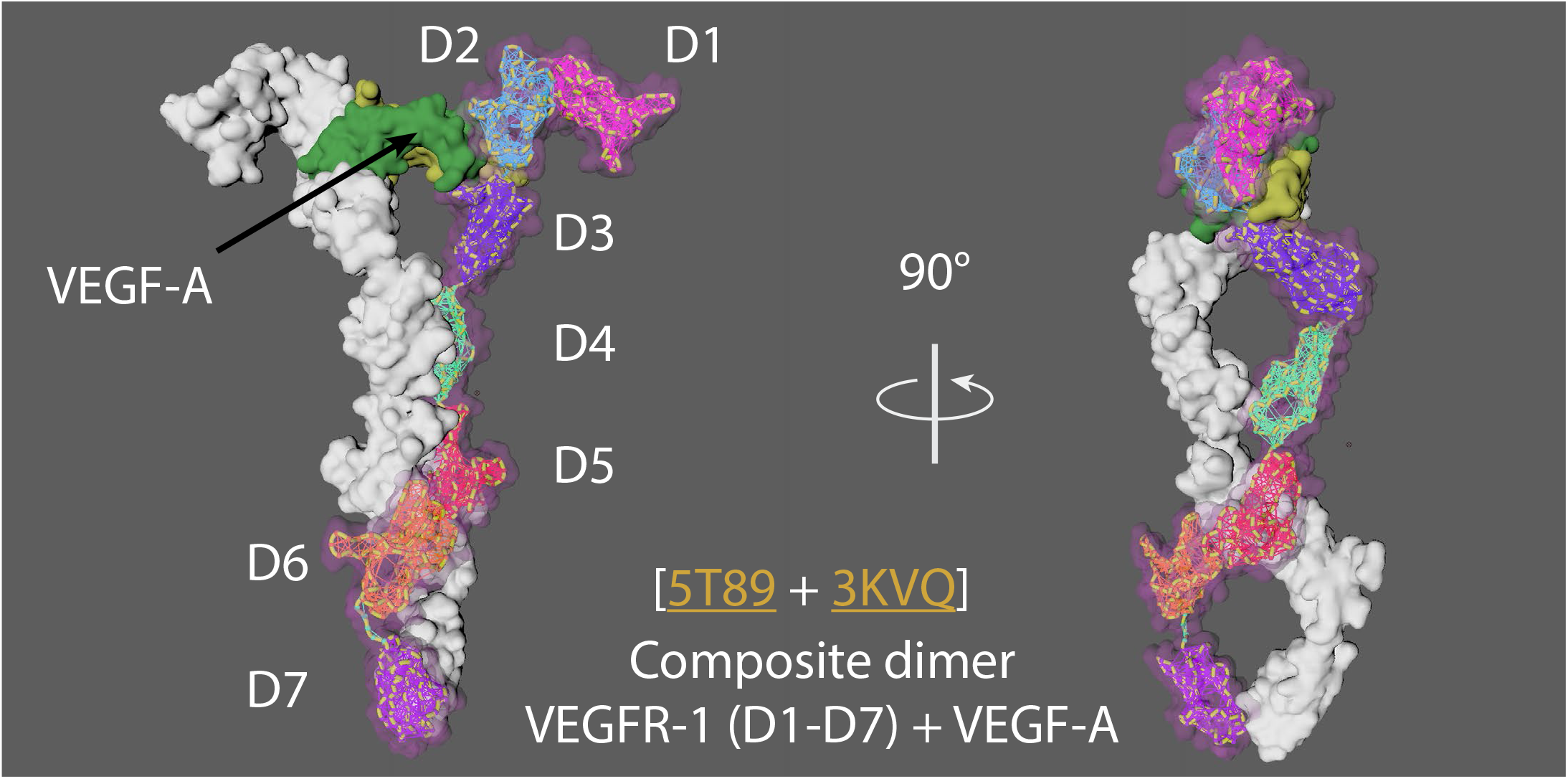

**MoA animation strategy:**

5Y89^1^ + 3KVQ2 = Composite VEGFR1 dimer1 from Markovic-Mueller et al. 2017; PDB available from Data S1.

The composite VEGFR1 dimer was rigged using mMaya rigging kit. Elastic networks were created for each domain (D1 to D7). Handles were made for the entire CA backbone and individual domains to manually pull the rig away from the dimer conformation, artistic licence was used to create a hypothetical monomer conformation (as no monomer structure is currently available). The hypothetical monomer was simulated to conformationally morph into the target composite VEGFR dimer for the cached animation.

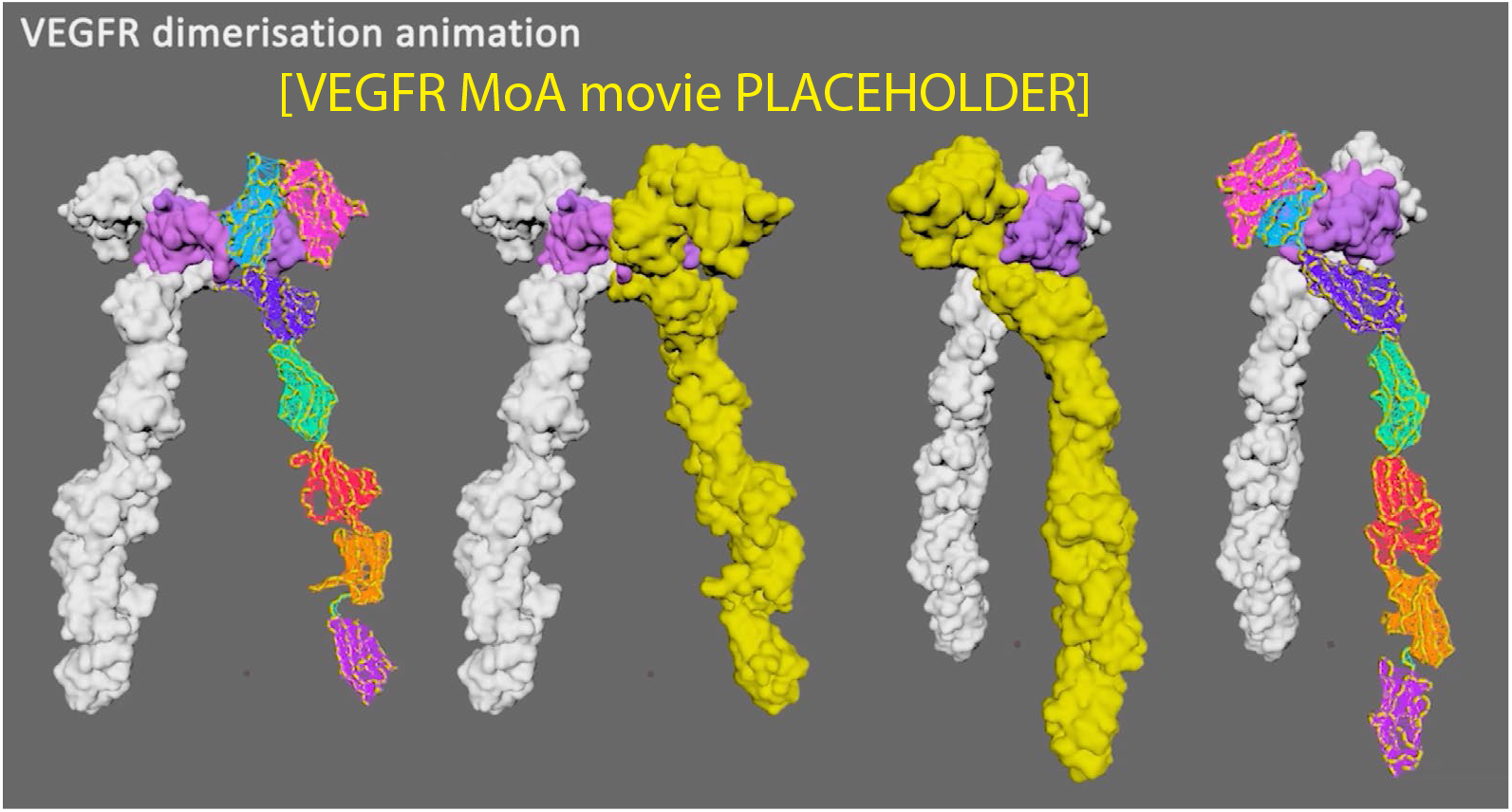

#### References

1. Markovic-Mueller, S., Stuttfeld, E., Asthana, M. et al. Structure of the Full-length VEGFR-1 Extracellular Domain in Complex with VEGF-A. Structure, 25 (2), 341-352, (2017). https://doi.org/10.1016/j.str.2016.12.012

2. Yang, Y., Xie, P., Opatowsky, Y. et al. Direct contacts between extracellular membrane-proximal domains are required for VEGF receptor activation and cell signalling. PNAS, 107 (5), 1906, (2010). https://doi.org/10.1073/pnas.0914052107

3. Sarabipour, S., Ballmer-Hofer, K. and Hristova K. VEGFR-2 conformational switch in response to ligand binding. Elife. 2016;5:e13876, (2016). https://doi.org/10.7554/eLife.13876

### c-KIT

- Stem Cell Factor (SCF) protomer binds directly to the D1, D2, and D3 ectodomain of c-KIT
- SCF binding leads to dimerisation of two c-KIT molecules
- This results in the lateral interaction of Ig-like domains D4 and D5 (i.e. D4 to D4 and D5 to D5) between the two monomers and brings the c-termini of the monomers 15Å of each other

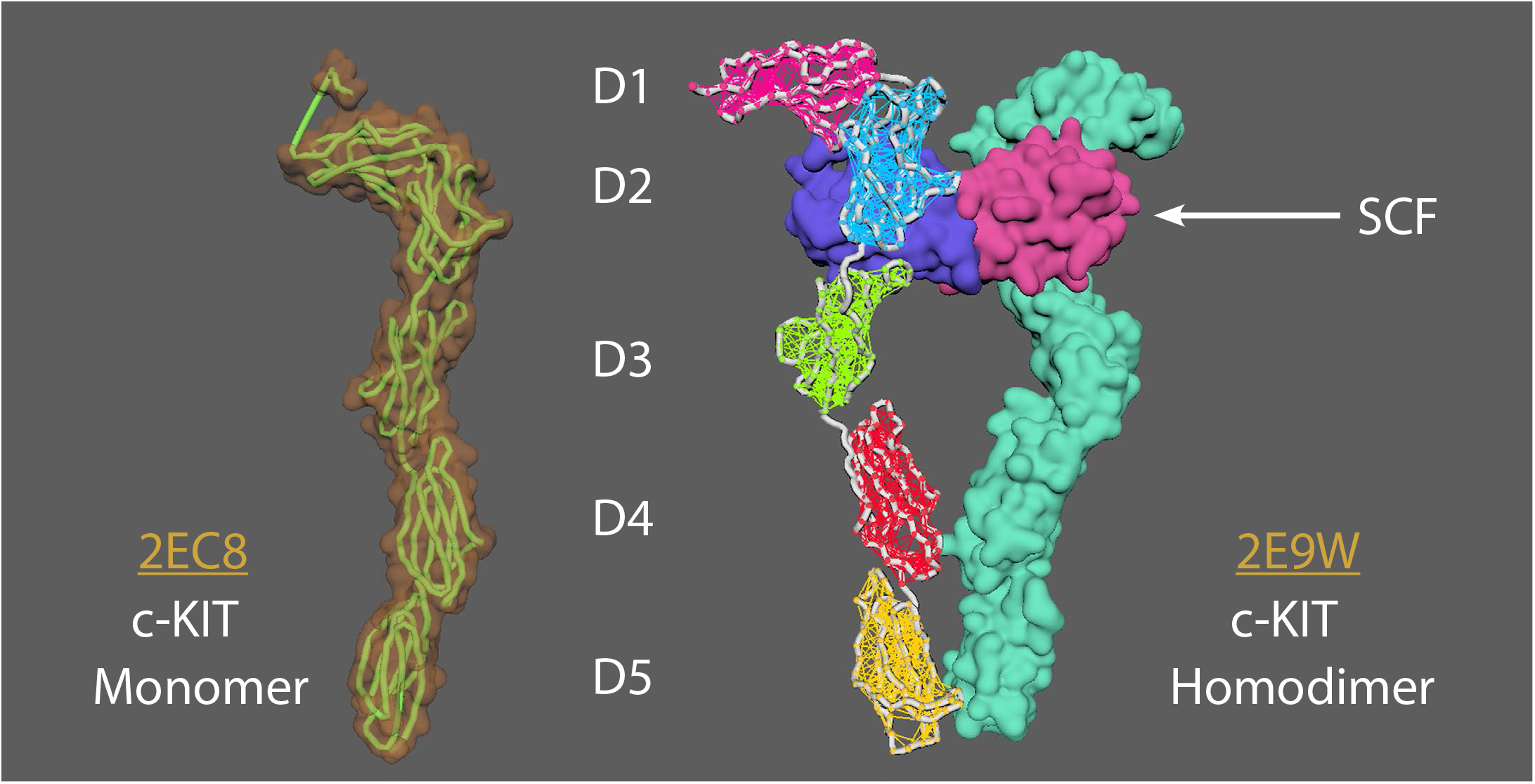

**MoA animation strategy:**

2EC8^1^ = c-KIT monomer

2E9W^1^ = c-KIT homodimer with SCF

Aligned chain A from 2E9W (dimer conformation) to 2EC8 (monomer conformation). Saved a new PDB version of chain A 2E9W (now as a monomer conformation). Created elastic networks for each domain (D1-5). Set it to target morph to the original 2E9W dimer position, cached animation.

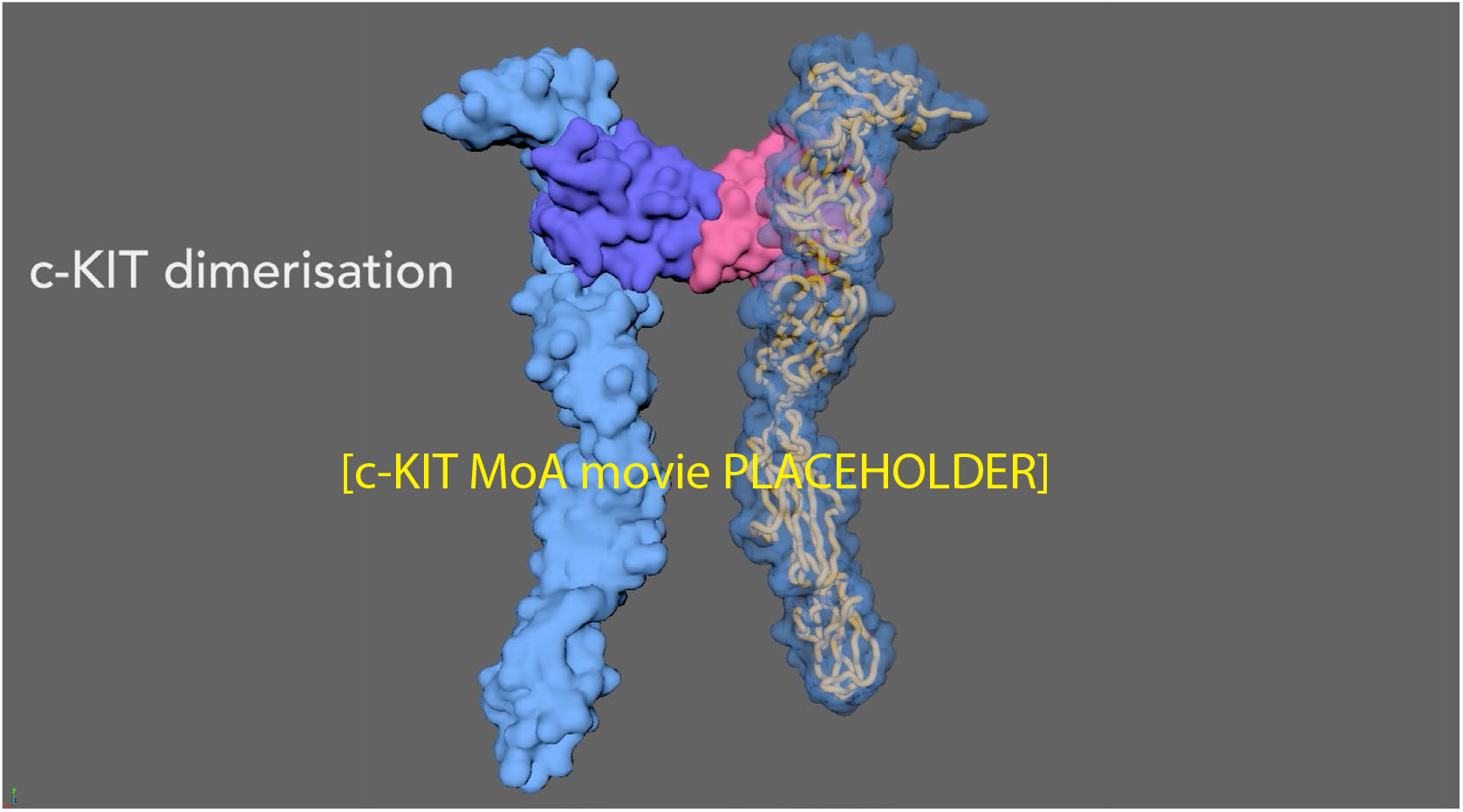

#### References

1. Yuzawa, S., Opatowsky, Y., Zhang, Z. et al. Structural Basis for Activation of the Receptor Tyrosine Kinase KIT by Stem Cell Factor. Cell, 130 (2), 323-334, (2007). https://doi.org/10.1016/j.cell.2007.05.055

2. Felix, J., De Munck, S., Verstraete, K. et al. Structure and Assembly Mechanism of the Signaling Complex Mediated by Human CSF-1 [published correction appears in Structure. 2020 Apr 7;28(4):488]. Structure. 23(9):1621-1631, (2015). https://doi.org/10.1016/j.str.2015.06.019

### Insulin Receptor (IR)

- IR is a dimer of heterodimers that comprises two α-chains and two β-chains, represented as (αβ)2.
- 2 insulin molecules per dimer. Each bound between L1 of one monomer and FnIII-1 of the other momoner.
- IR goes from an inverted U-shaped dimer to a T-shaped dimer upon ligand binding. One ligand binding to one receptor can lead to the conformational change.

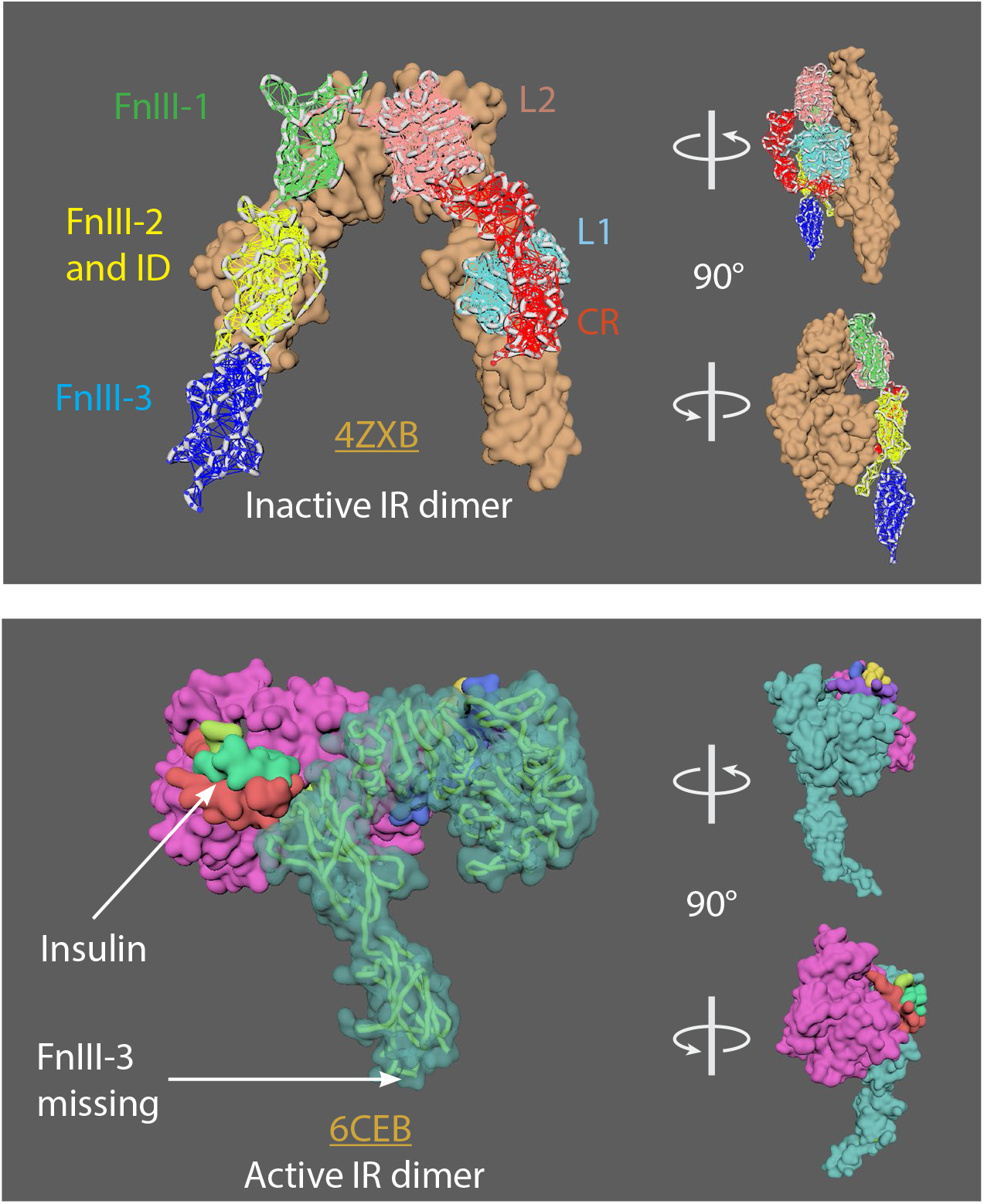

**MoA animation strategy:**

4ZXB^1^ = inactive model (-ligands) 6CEB^2^ = active model (+ ligands)

The missing FnIII-3 domain in 6CEB chain A was added using the mMaya modelling kit to get the whole ectodomain structure (FnIII-3 domain taken from 4ZXB); a new PDB was created. A mMaya rig was made for 4ZXB and elastic networks created for L1, CR, L2, FnIII-1, FnIII-2 and ID, and FnIII-3 domains. Rig was target morphed to the new version of 6CEB, mesh animation was cached. Binding animation of insulin ligands on 4ZXB were keyframed manually and was approximated based on their position relative to 6CEB.

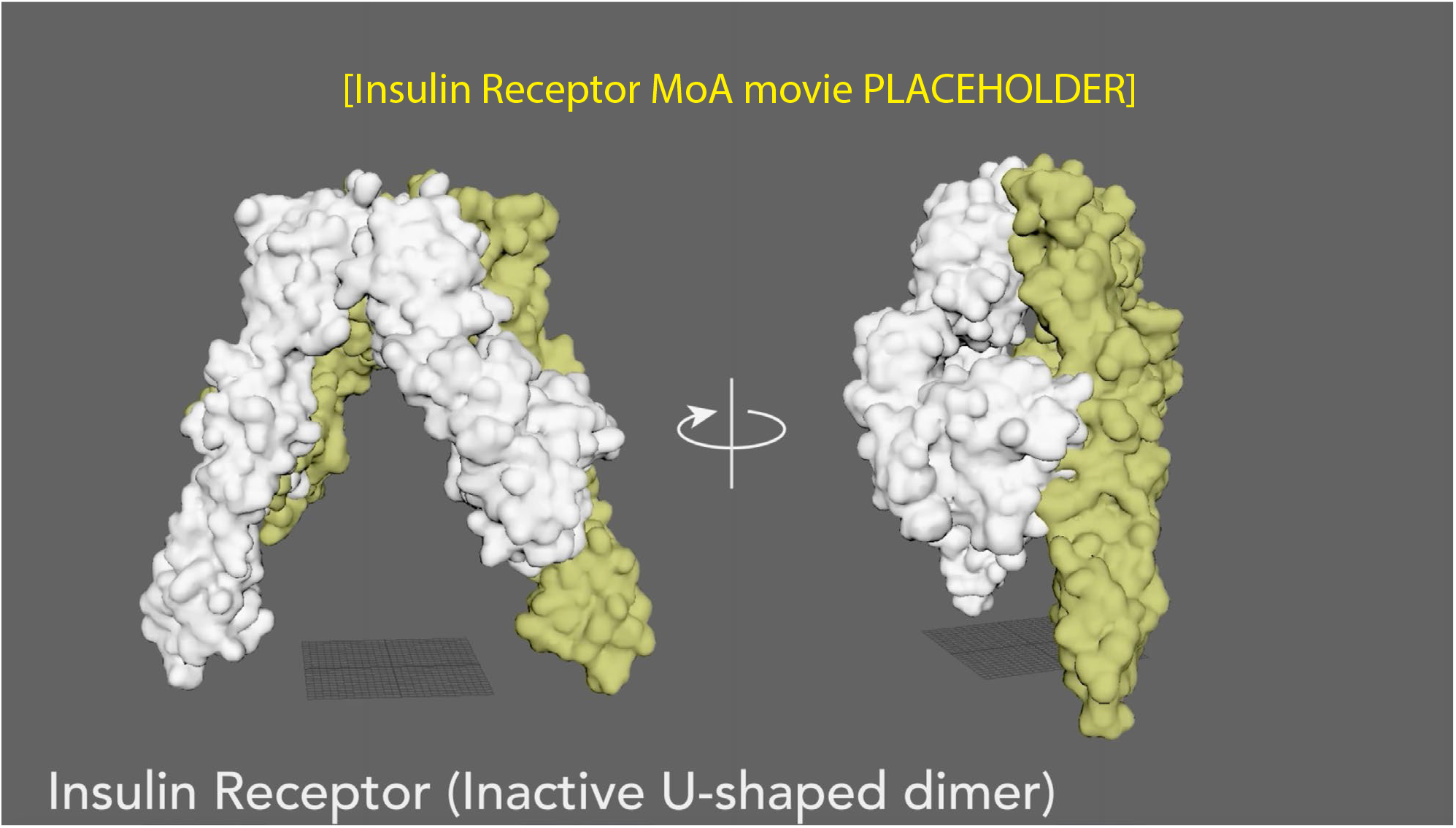

#### References

1. Croll, T., Smith, B., Margetts, M. et al. Higher-Resolution Structure of the Human Insulin Receptor Ectodomain: Multi-Modal Inclusion of the Insert Domain. Structure, 24 (3), 469-476. (2016) https://doi.org/10.1016/j.str.2015.12.014

2. Scapin, G., Dandey, V., Zhang, Z. et al. Structure of the insulin receptor–insulin complex by single-particle cryo-EM analysis. Nature 556, 122–125 (2018). https://doi.org/10.1038/nature26153

3. Gutmann, T., Kim, K., Grzybek, M. et al. Visualization of ligand-induced transmembrane signaling in the full-length human insulin receptor. J Cell Biol 7; 217 (5): 1643–1649 (2018) https://doi.org/10.1083/jcb.201711047

### Tetraspanin CD81 (TAPA-1)

- Tetraspanin CD81 structure resembles a waffle cone when bound with cholesterol
- Cholesterol binding regulates CD81-mediated export of CD19
- EC2 extracellular domain covers an intramembrane cavity of 4 transmembrane helices (TM 1-4)
- In the absence of cholesterol, EC2 adopts an ‘‘open’’ conformation
- A salt bridge between EC2 and TM4 (D196 – K201) stabilises the closed conformation and breaks during opening.
- Another salt bridge (K116 – D117) forms upon opening which stabilises the open conformation

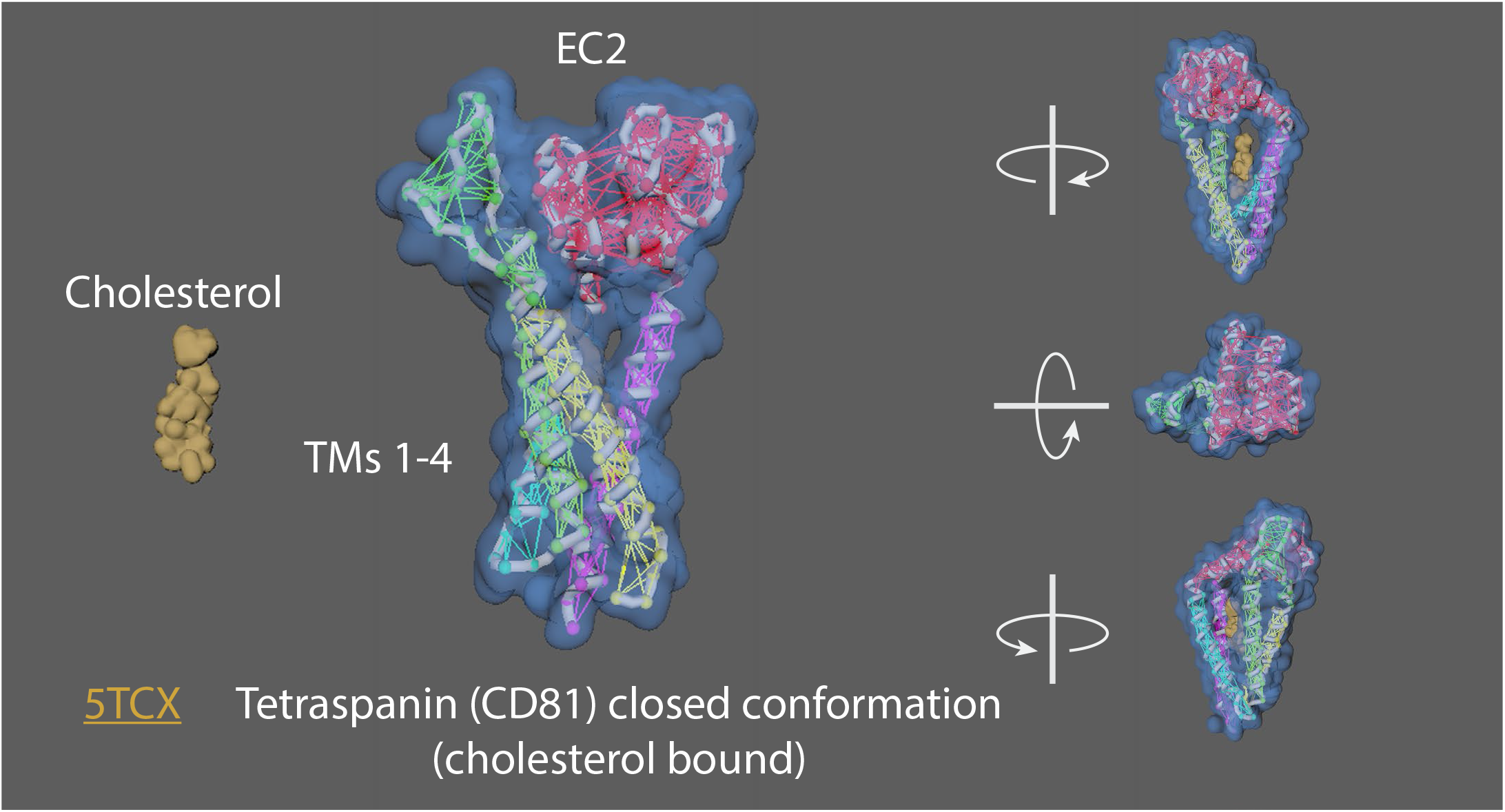

**MoA animation strategy:**

5TCX^1^ = Tetraspanin (closed conformation i.e. cholesterol-bound)

Rigged 5TCX, created elastic networks for EC2 (residues 34-55 and 113-201), TM1 (6-33), TM2 (56-84), TM3 (86-112), TM4 (202-232). Created a handle for the residues 113-201 of the EC2 extracellular domain, manually keyframed to form the open conformation (apoprotein/unbound to cholesterol).

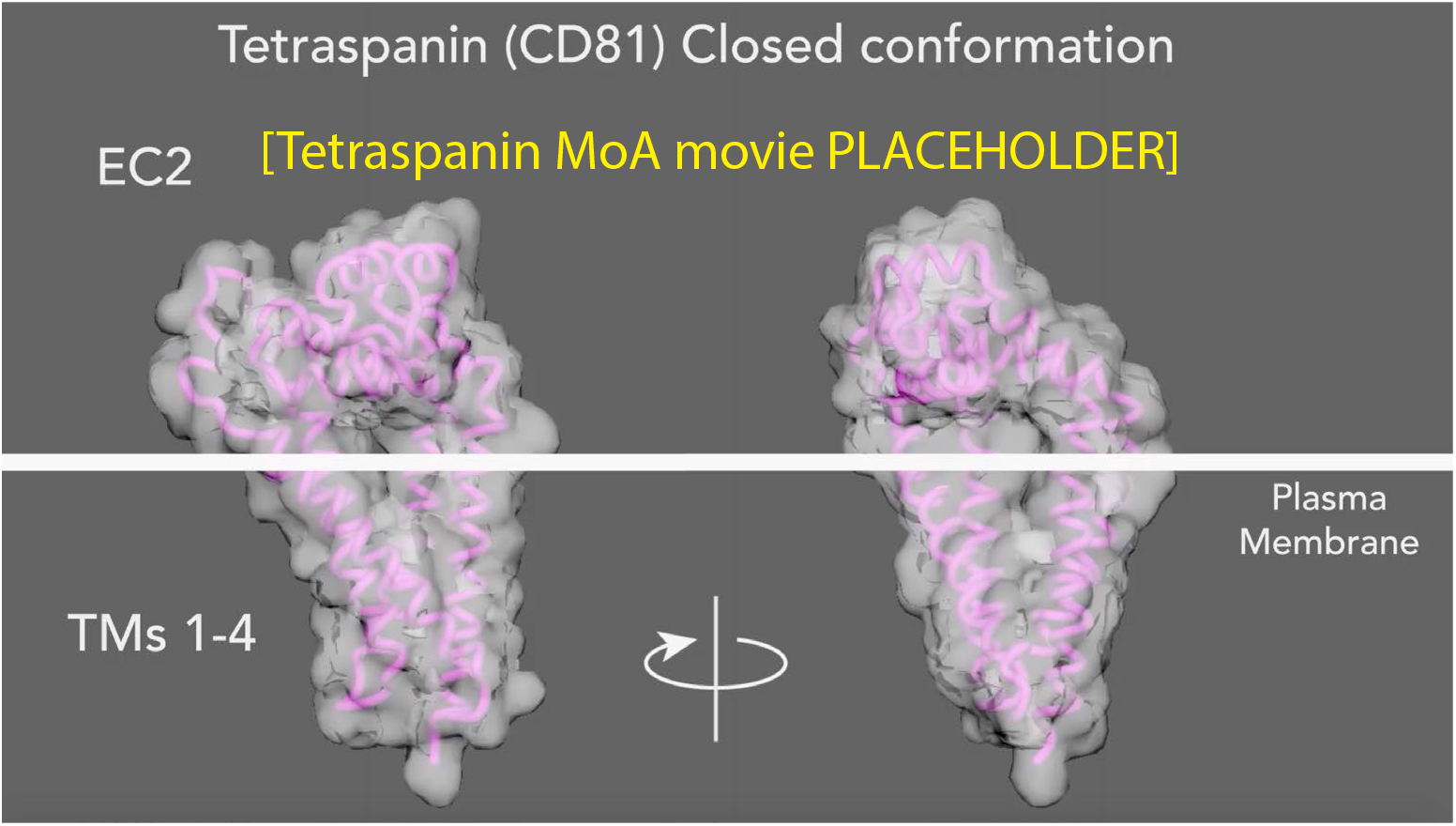

#### References

1. Zimmerman, B., Kelly, B., McMillan, B. et al. Crystal Structure of a Full-Length Human Tetraspanin Reveals a Cholesterol-Binding Pocket. Cell, 167 (4), 1041-1051.e11, (2016). https://doi.org/10.1016/j.cell.2016.09.056

### TNFR super family

- In the TNFR superfamily there are 3 groups of receptors: 1) Death receptors (DRs); 2) TNFR-associated factor (TRAF)– interacting receptors, and 3) decoy receptors (DcRs)
- Basic unit of signalling is a trimeric ligand and three receptors.
- Each receptor binds on the outside interface of 2 ligand monomers
- Receptors can pre-assemble (as resting or “nonsignaling” state) on the cell surface and can form either parallel or antiparallel dimers.
- Preferred model is antiparallel dimers
- Antiparallel dimers are formed between the receptor CRD1 and CRD2 domains which occludes the ligand-binding site.
- Arranged as a large hexagonal lattice where each point connects 3 receptor monomers.
- Individual ligand-receptor complexes ~120 Å apart. Total edge length of ~170 Å (may vary with receptor type).
- Ligand binding leads to a conformational change in receptors (into an upright position, perpendicular to the cell surface) allowing cell signaling but maintaining hexagonal symmetry.

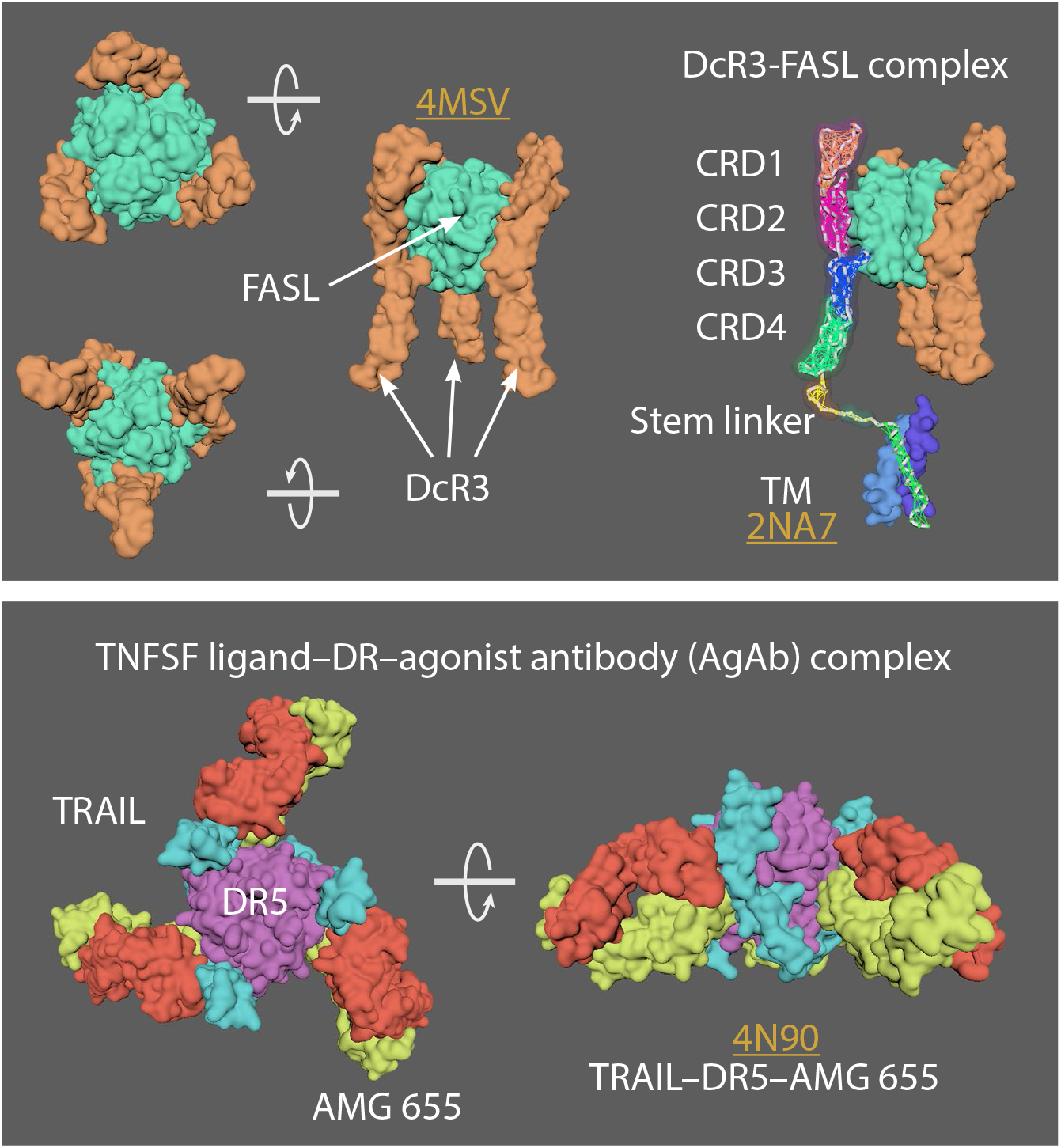

**MoA animation strategy:**

4MSV^1^= Decoy receptor 3 (DcR3)(chain A) and FasL (chain B)

2NA7^2^ = Transmembrane domain of human Fas/CD95 death receptor

4N90^3^ = Crystal structure of TRAIL-DR5 with the agonist antibody

As no complete structures of the active state or resting/non-signalling state structures exist, a “hybrid” TNFRSF molecule was made. Created a stem linker (PQIEN VKGTE DSGTT) taken from TNFR1 (1EXT4, residues 197-211) with mMaya modelling kit to connect the CRD4 of DcR3 (4MSV) and a TM domain trimer (2NA7). New PDB created named “Hybrid TNFRSF” and rigged with mMaya rigging kit. Elastic networks of CRD1-4, stem linker and TM domain were created. Handles were created (CD1+2; CD3, and CD4) to manually move the rig into a hypothetical resting state. Movement was keyframed, animation cached and meshes extracted. Alembic cached meshes were positioned in a hexagonal lattice and ligand binding was keyframed for the final animation. 4N90 [Crystal structure of TRAIL-DR5 with the agonist antibody (Fab fragments)] was used as a reference to get the correct spacing of the hexagonal array when the trimers become active (from their antiparallel dimer conformation in the resting state).

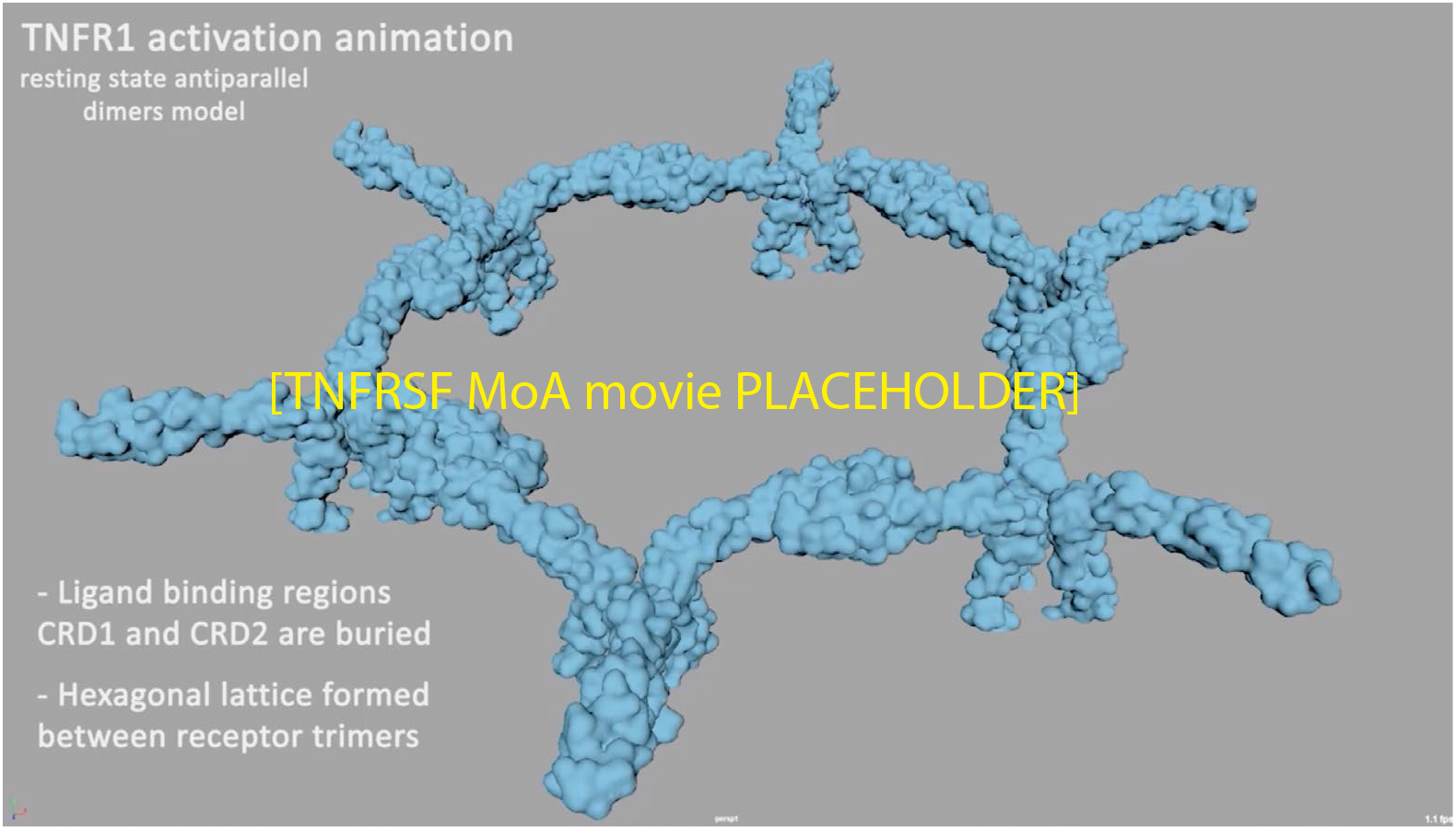

#### References

1. Liu, W., Ramagopal, U., Cheng, H. et al. Crystal Structure of the Complex of Human FasL and Its Decoy Receptor DcR3. Structure, 24 (11), 2016-2023, (2016). https://doi.org/10.1016/j.str.2016.09.009

2. Fu, Q., Fu, T., Cruz, A. et al. Structural Basis and Functional Role of Intramembrane Trimerization of the Fas/CD95 Death Receptor. Molecular Cell, 61 (4), 602-613, (2016). https://doi.org/10.1016/j.molcel.2016.01.009

3. Graves, J., Kordich, J., Huang, T. et al. Apo2L/TRAIL and the Death Receptor 5 Agonist Antibody AMG 655 Cooperate to Promote Receptor Clustering and Antitumor Activity. Cancer Cell, 26 (2), 177-189, (2014). https://doi.org/10.1016/j.ccr.2014.04.028

4. Naismith, J., Devine, T., Kohno, T. et al. Structures of the extracellular domain of the type I tumor necrosis factor receptor. Structure, 4 (11), 1251-1262, (1996). https://doi.org/10.1016/S0969-2126(96)00134-7

5. Vanamee, É. and Faustman, D. Structural principles of tumor necrosis factor superfamily signaling. Science Signaling, 11 (511), eaao4910, (2018) https://doi.org/10.1126/scisignal.aao4910

### GLUT1

- GLUT1 (glucose transporter 1) is over-expressed in many cancer cells
- 12 transmembrane (TM) segments form the N-terminal and C-terminal domains
- Rotation of the N-terminal and C-terminal domains occurs to transition from an outward-open to an outward-occluded, and finally an inward-open state to allow D-glucose transport

**MoA animation strategy:**

4PYP^1^ = GLUT1; inward-open state

4ZWC^2^ = GLUT3; outward-open state

Rigged 4PYP (chain A) (inward-open state) was targetted to conformationally morph ‘backwards’ into 4ZWC (outward-open state), and the alembic cache was reversed. Created elastic networks of N-terminal domain (9-206), C-terminal domain(272-455), and ICH domain (211-264).

Downloaded Het atoms for D-glucose structure and manually keyframed glucose uptake.

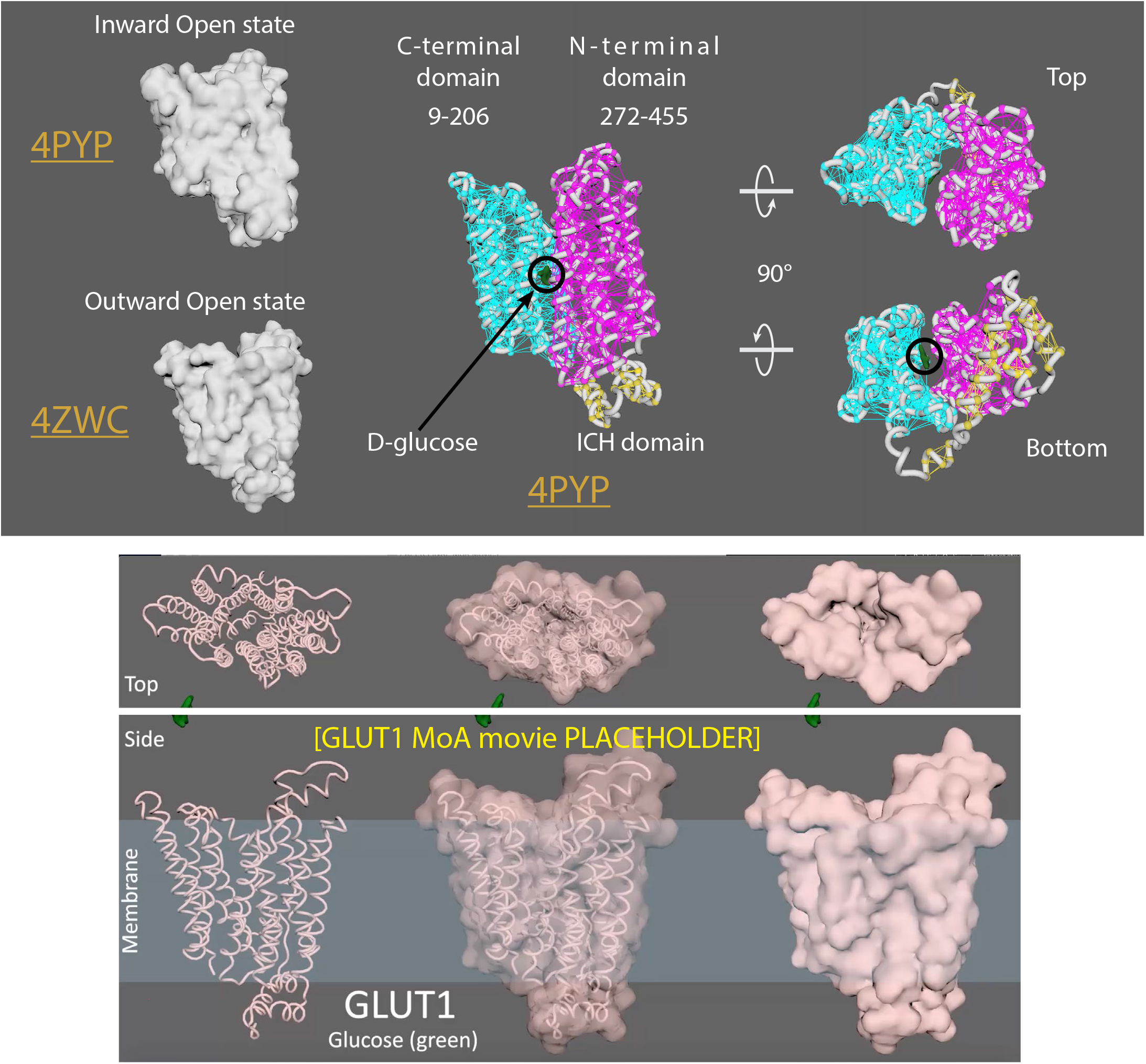

#### References

1. Deng, D., Sun, P., Yan, C. et al. Molecular basis of ligand recognition and transport by glucose transporters. Nature 526, 391–396 (2015). https://doi.org/10.1038/nature14655

2. Deng, D., Xu, C., Sun, P. et al. Crystal structure of the human glucose transporter GLUT1. Nature 510, 121–125 (2014). https://doi.org/10.1038/nature13306

